# Comparative genomics provides an operational classification system and reveals early emergence and biased spatio-temporal distribution of SARS-CoV-2

**DOI:** 10.1101/2020.06.26.172924

**Authors:** Matteo Chiara, David S. Horner, Carmela Gissi, Graziano Pesole

## Abstract

Effective systems for the analysis of molecular data are of fundamental importance for real-time monitoring of the spread of infectious diseases and the study of pathogen evolution. While the Nextstrain and GISAID portals offer widely used systems for the classification of SARS-CoV-2 genomes, both present relevant limitations. Here we propose a highly reproducible method for the systematic classification of SARS-CoV-2 viral types. To demonstrate the validity of our approach, we conduct an extensive comparative genomic analysis of more than 20,000 SARS-CoV-2 genomes. Our classification system delineates 12 clusters and 4 super-clusters in SARS-CoV-2, with a highly biased spatio-temporal distribution worldwide, and provides important observations concerning the evolutionary processes associated with the emergence of novel viral types. Based on the estimates of SARS-CoV-2 evolutionary rate and genetic distances of genomes of the early pandemic phase, we infer that SARS-CoV-2 could have been circulating in humans since August-November 2019. The observed pattern of genomic variability is remarkably similar between all clusters and super-clusters, being UTRs and the s2m element, a highly conserved secondary structure element, the most variable genomic regions. While several polymorphic sites that are specific to one or more clusters were predicted to be under positive or negative selection, overall, our analyses also suggest that the emergence of novel genome types is unlikely to be driven by widespread convergent evolution and independent fixation of advantageous substitutions. While, in the absence of rigorous experimental validation, several questions concerning the evolutionary processes and the phenotypic characteristics (increased/decreased virulence) remain open, we believe that the approach outlined in this study can be of relevance for the tracking and functional characterization of different types of SARS-CoV-2 genomes.

## Introduction

The ongoing Coronavirus disease 2019 (COVID-19) pandemic [1] poses the greatest global health and socioeconomic threat since world war II. The first case of COVID-19 was reported in Wuhan city, Hubei province, China, in late December 2019 [2], although retrospective analyses have placed the onset as early as December 1st.

The most common symptoms include fever, dry-cough and a general sense of fatigue, while some patients also experience muscle pain, nasal congestion, runny nose, sore throat, or diarrhoea [3-4]. In a minority of the patients the infection may cause pneumonia, severe acute respiratory syndrome, kidney failure and even death [5-6]. Symptoms are usually mild in children and young adults, while elderly, immunosuppressed, and individuals affected by a cardiac disease or diabetes are at higher risk of developing severe symptoms [7].

COVID-19 is primarily transmitted between people through respiratory droplets and contact routes [8], although some recent studies suggest that airborne transmission might also be possible [9-10]. The incubation period typically ranges between 2 and 14 days, although longer incubation times have been also reported [11]. Notwithstanding the development of several promising therapeutic approaches [12], at present no universally approved therapeutic strategy is available for the treatment of COVID-19.

At the time of writing, COVID-19 has affected more than 200 countries worldwide, with more than 9 Million confirmed individual infections and a death toll in excess of 480 thousand. The limited availability of diagnostic kits in several countries, varying criteria for reporting COVID-19-related deaths, and the fact that very mild or asymptomatic infections can often go undetected [13], suggest that both these figures are likely to represent substantial underestimates of the worldwide impact of SARS-CoV-2. The first complete genomic sequences of the viral pathogen were determined in early January 2020 by Next Generation Sequencing metatranscriptomics [14], allowing the rapid development of diagnostic tests (Corman et al., 2020) and the development of molecular monitoring strategies [15-16].

The genome is approximately 30.000 nt in size and shows high similarity (∼79%) with SARS-CoV [17], a beta-coronavirus of the subgenus Sarbecovirus, and the causal agent of a large scale epidemic of viral pneumonia (Severe Acute Respiratory Syndrome, SARS) that hit China and other 25 countries in 2003 and 2004 [18]. The International Committee on Taxonomy of Viruses (ICTV) has designated the novel pathogen as SARS-CoV-2. Phylogenetic analyses of conserved protein domains of more than 2500 coronaviruses have been used to assign SARS-CoV-2 to the group of the Severe acute respiratory syndrome-related coronavirus species (SARSr-CoV), where it forms a relatively distant sister group to SARS-CoV, interleaved with various SARSr-CoV isolated from non-human mammalian species [19].

SARS-CoV-2 shows the highest levels of genome identity (96%) with a bat SARS-related (SARS-r) CoV denoted RaTG13, isolated in the Yunnan province [20]. Despite this sequence similarity, SARS-CoV-2 differs from RaTG13 in several key features. Arguably, the most important is the presence of a polybasic furin cleavage site insertion (residues PRRA) at the junction of the S1 and S2 subunits of the Spike protein [21]. This insertion, which may increase the infectivity of the virus, is not present in related beta-coronaviruses, although similar polybasic insertions are observed in other human coronaviruses, including HCoV-HKU1, as well as in highly pathogenic strains of avian influenza virus [22]. Additionally, the RBD (Recognition Binding Domain) of the SARS-CoV-2 spike protein is significantly more similar (97% identity) to that of SARSr-CoVs isolated from specimens of Malayan pangolins (*Manis javanica*) illegally imported into southern China (Guangdong and Guangxi provinces) than to the RDB of RaTG13 (89% identity). This data suggest the possibility that pangolins or other mammalian species could have acted as “intermediate” or “amplifying” hosts for human transmission of SARS-CoV-2 [23-24].

As the COVID-19 pandemic has progressed, the number of viral isolates for which a genomic sequence is available has increased substantially. Currently, in the excess of 50.000 viral genomes are publicly available in dedicated repositories [25]. As expected, considering the recent ancestry, and reportedly low mutation rates of coronaviruses [26], SARS-CoV-2 genomes are highly similar (average identity 99.99%) and show reduced genetic diversity. This low diversity notwithstanding, the availability of a considerable number of genomic sequences, with worldwide sampling, allows the identification of phylogenetically distinct clusters of SARS-CoV-2 sequences [27-32]. Interestingly, several of these clusters show highly biased geographic distributions, and in many independent studies, the genomic signatures of different clusters have been tentatively linked to increased/decreased virulence or possible adaptation to human hosts [33-35].

While it should be stressed that, in the absence of careful experimental validation, it is extremely difficult to determine if the identified SARS-CoV-2 genetic variants have increased/decreased virulence and reflect adaptive evolution or genetic drift and founder effects, the importance of establishing a simple and reproducible system for the delineation of genomic diversity of human pathogens is universally acknowledged [36-37]. Indeed such a classification system could be extremely useful for the rapid identification of new emerging types and, more importantly, for a fine grained monitoring of the diffusion of pathogens in different geographic contexts over time.

Currently, the most widely used models for the classification of SARS-CoV-2 viral types are those provided by the curators of the GISAID [38] and Netxstrain portals [39], which incorporate the most complete and accurate resources for comparative genomics of SARS-CoV-2, Providing extensive databases of more than 40000 viral sequences with associated metadata, including date and place of isolation, as well as information concerning health and background data of the patients. GISAID also provides tools for comparative genomic and phylogenetic analyses to facilitate the annotation of the genomes, their classification in types/groups and the functional interpretation of the data. Similarly, Nexstrain collects a selection of publicly available genomes and associated metadata, to provide informative analyses and graphical representation of the spread of SARS-CoV-2 over time as well as detailed and curated phylogenetic analyses for the classification of genome types. Although both platforms constitute invaluable resources for the SARS-CoV-2 research community, the associated classification systems suffer from some inherent limitations. The most important is that the criteria used to establish the clusters/types of SARS-CoV-2 are not set out clearly, and that the delineation of clusters is based on arbitrary manual annotation of phylogenetic trees, rather than on inherent genomic features of SARS-CoV-2, limiting the reproducibility of results. Furthermore, Nextstrain provides classification only for a restricted selection of the available viral genomes (currently less than 4000). Finally, limited variability and the recurrent events of recombination observed in Sarbecovirus [40-41] may hinder the validity of classification systems based purely on phylogenetic analyses and manual annotation.

In the light of these considerations, in the present work we propose a simple set of rules, which could serve as an operational classification system of SARS-CoV-2 genomic sequences. The proposed classification system is inspired by MultiLocus Strain Typing (MLST), a classification approach normally used for microbes [42]. Thus, for a given dataset of available SARS-CoV-2 genomes, it combines the empirical study of genomic variability with the analyses of the high frequency variant sites that are fixed in the extant viral population. In this way our approach derives a simple but effective set of rules for a systematic and highly reproducible delineation of SARS-CoV-2 genomes, that are therefore grouped in clusters and superclusters.

To demonstrate the validity of our approach, we conducted an extensive comparative genomic analysis of more than 20,000 SARS-CoV-2 complete genomes. By applying the proposed classification system we derived important observations concerning different types of SARS-CoV-2, their evolutionary pattern and geographic distribution, and the mechanisms associated with the emergence of possible novel types.

## Materials and methods

A collection of 20,521 putatively complete high coverage SARS-CoV-2 genomes and associated metadata was retrieved from the GISAID EpicoV [38] platform on 28th May 2020. A total of 13 SARSr-CoV genomic sequences isolated from non-human hosts, including bats and pangolins [23-24], were also retrieved from the GISAID EpiCoV portal at the same date. SARS-CoV-2 sequence comparisons were performed using the reference Refseq [43] assembly NC_045512.2, collected on 26th December 2019 and identical to the sequence of the oldest SARS-CoV-2 isolates, dating back to 24th December 2019 (EPI_ISL_402123).

SARS-CoV-2 genomes were aligned to the 29,903 nt-long reference assembly of SARS-CoV-2 by means of the *nucmer* program [44]. Custom Perl scripts were used to infer the size of each genomic assembly and the number of uncalled bases/gaps (denoted by N in the genomic sequence). Then, we analysed only the 11,633 high quality complete genomes, defined as those longer than 29,850 nt and including less than 150 ambiguous sites.

Variant sites, including substitutions and small insertion and deletions, were identified by using the *show-snps* utility of the *nucmer* package. Output files were processed by the means of a custom Perl script, and converted into a phenetic matrix, with variable positions on the rows and viral isolates in the columns. Values of 1 and 0 were respectively used to indicate presence or absence of a variant.

Genetic distances between genomic sequences were established from this phenetic matrix using the *dist* function of the R *stat* package with default parameters (Euclidean distances) [45-46]. Clusters were established by means of hierarchical clustering algorithms, with complete linkage as implemented in the *hclust* R standard libraries function. The *cutree* function was used to separate distinct clusters at the desired level of divergence (2 distinct variant sites).

Functional effects of genetic variants, as identified from genome alignments, were predicted by means of a custom Perl script, based on the annotation of the NC_045512.2 SARS-CoV-2 reference assembly.

Identification of sites possibly under selection was performed by applying the MEME and FEL methods, as implemented in the *Hyphy* package[47], on the concatenated alignment of protein coding sequences of all the 11,633 previously identified high quality complete SARS-CoV-2 genomes. A p-value of 0.05 was considered for the significance threshold.

A total of 68 viral genomes of the SARS 2003 outbreak were retrieved from the NCBI virus database [48]. Classification/association of strains to the 3 (early/middle/late) phases of the SARS 2003 epidemic are according to Song et al 2005 [49].

Calculation of evolutionary rates of SARS-CoV-2 and estimation of times of divergence were performed according to the formula described in Zhao et al [50], based on genetic distances as determined in this study.

Analyses of prevalence of allele frequency over time were executed based on the collection dates of individual genomes as reported in the GISAID metadata table. The collection date of the reference genomic sequence of SARS-CoV-2 in GISAID (26th December 2020), was set as time 0. Consecutive, non-overlapped intervals of 10 by 10 days were considered. A total of 1381 genomic sequences, for which collection dates were not reported in GISAID, were excluded from these analyses.

Comparison of levels of variability of “early” and “late” clusters of SARS-CoV-2 genomes were established by 100 random resampling of 150 genomes (batch), matched by date of collection, from each defined cluster (see below). The total number of distinct variant sites was calculated for each random batch of genomes, in order to derive a distribution of genomic variability. Significance between distributions of genomic diversity were established by means of the Wilcoxon, Sum and Rank test as implemented in the standard R libraries [45].

Variability with respect to the reference NC_045512.2 SARS-CoV-2 genome was computed on sliding windows of 100 bp, overlapped by 50 bp, by counting the proportion of variable sites contained in each window (number of variable sites in the window, divided by the total number of variable sites in the entire genome) with a custom Perl script. A Fisher-exact test, contrasting the local variability in a window with the average variability in the genome, was used to identify hypervariable regions. P-values were corrected using the Benjamini-Hochberg procedure for the control of False Discovery Rate.

Predictions of the secondary structure of the “Coronavirus stem-loop II-like motif” (s2m) and its Minimum Folding Energy (MFE) calculation were estimated with the RNAfold program [51] of the Vienna package [52], by artificially implanting each of the possible 129 substitutions in the 43 nt-long s2m sequence identified in the reference SARS-CoV-2 genome and in the presumably ancestral sequence of s2m, as observed in the genome of the RatG13 SARSr-CoV-2.

Prediction of the consensus co-folding structure of s2m in SARS-CoV-2 was obtained by applying the R-scape[53] program to the alignment of all s2m sequences found in the collection of the 11,633 high quality complete genome analysed in this study.

Consensus secondary structure of the s2m element of Coronaviruses was as in the model RF00164 (https://rfam.org/family/RF00164) of the RFAM database (reference).

Graphical representation of the data, basic statistical analyses and clustering of viral genomes were performed by means of the standard libraries of the R programming language.

The software for the operational classification of SARS-CoV-2 lineages proposed here, is publicly available through this github repository https://github.com/matteo14c/assign_CL_SARS-CoV-2.

## Results

### Genomic features and evolutionary dynamics of SARS-CoV-2

We retrieved 20,521 SARS-CoV-2 genomic sequences labeled as high coverage and putatively complete, covering 86 countries in 5 continents from the GISAID EpiCoV portal. Surprisingly, we observed that a considerable number (6462) of the reportedly complete genomic assemblies of SARS-CoV-2 presented incomplete 3’ or 5’ UTRs (median size 29782 nts; reference genome size of 29,903 nts, including a polyA tail of 33 nts). Additionally, 4310 genomes contained a large number of gaps and/or uncalled bases - ranging from 151 to 3500. Stringent criteria were used to retain only sequences which were more likely to represent a nearly complete assembly of the SARS-CoV-2 genome (more than 29,850 nts in size) and contained only a limited number of missing positions (less than 150 Ns). A total of 11,633 genomes were therefore selected and analysed (Supplementary Table S1).

Comparative analysis of genetic distances between these SARS-CoV-2 genomes (see Materials and methods) showed an average number of 0.64 variant sites per nearest neighbor pair. The equivalent figure for late-phase isolates of SARS-CoV from the SARS 2003-2004 epidemics [49] was slightly higher (0.78 polymorphic sites per nearest neighbor pair). Consistent with these observations, estimates of mutation rates, according to the formula described in Zhao et al [50] are marginally lower for SARS-CoV-2 (1.84 x 10^−3^ substitutions per site per year) compared to SARS-CoV (2.38 x 10^−3^ substitutions per site per year). Interestingly, we notice that 6384 (54.87%) of the high quality SARS-CoV-2 genomes here analysed have a perfect sequence identity (0 polymorphic sites) with respect to their closest neighbor. All in all these observations are consistent with the reportedly low mutation rate of coronaviruses with respect to other single strand RNA viruses [26,50].

A total of 6319 distinct variant sites were observed between the 11,633 high quality genomes included in these analyses (Supplementary Table S2). 99.22% of these sites have an allele frequency below 1%, the threshold above which a variant is often considered fixed in a natural population [54]. This value is not surprising, consideringly the relatively recent ancestry and the low mutation rate of SARS-CoV-2. Thus, only 50 variant positions (0.88%) show an allele frequency of 1% or greater (Table 1).

**Table 1:**
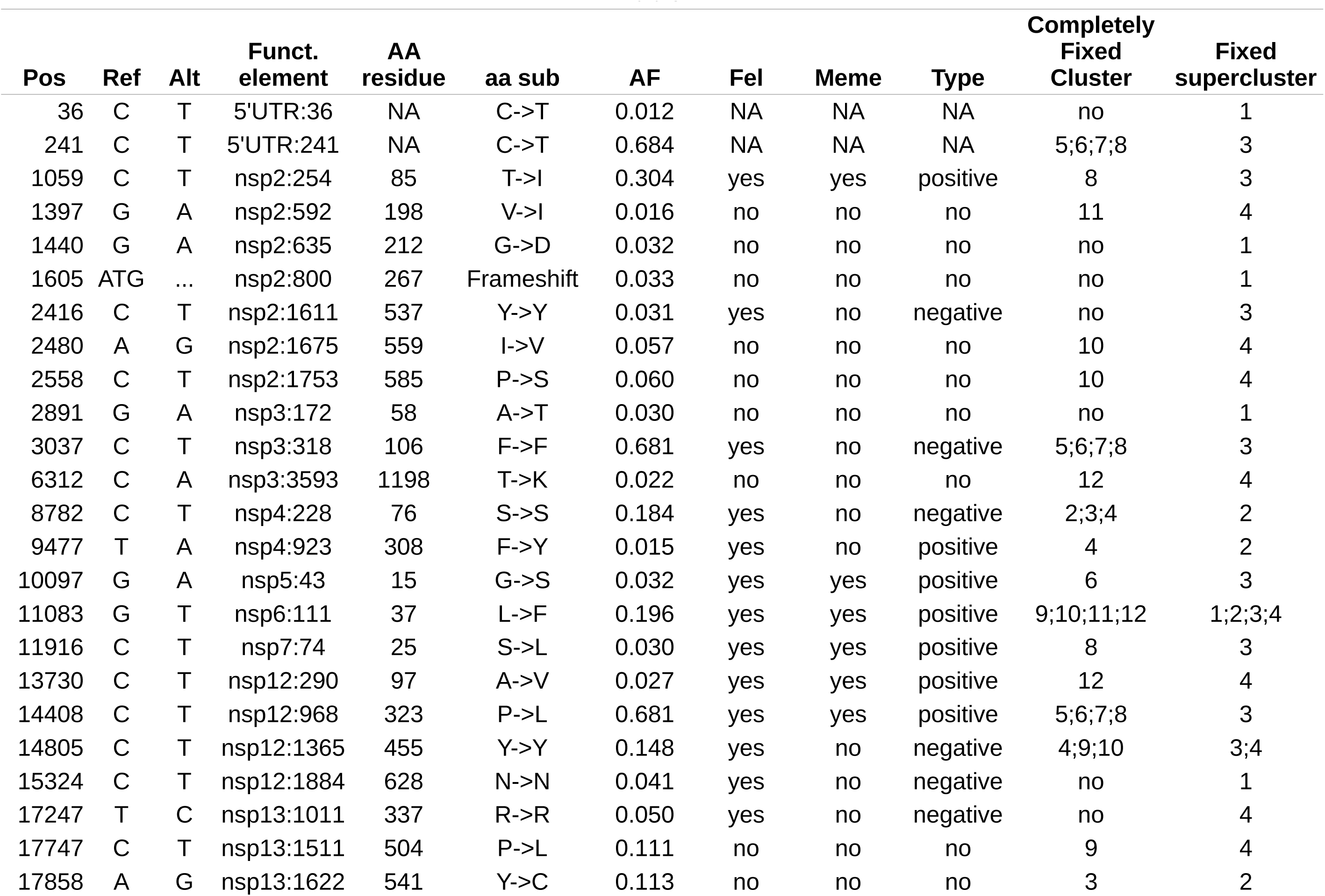
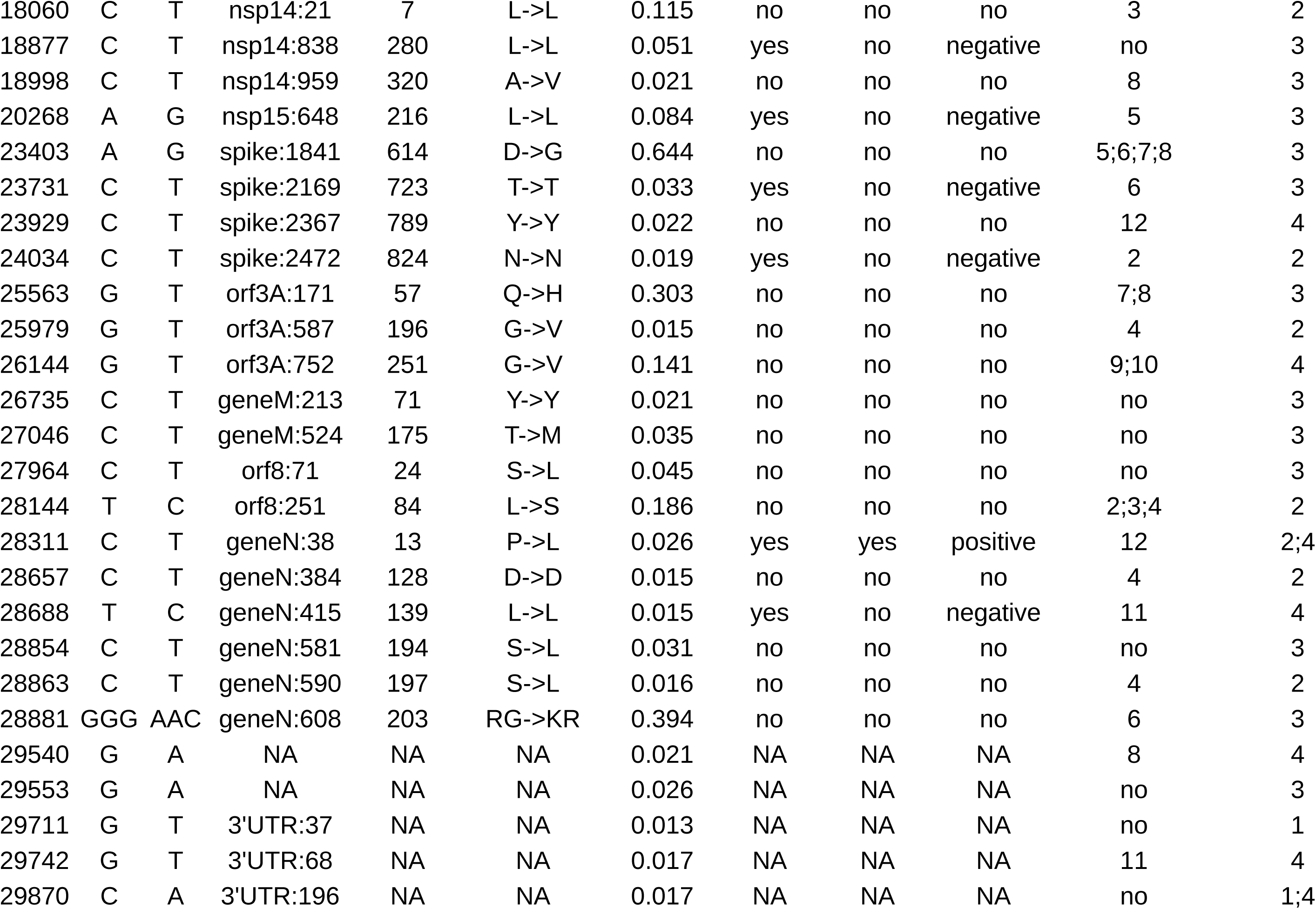
List of high frequency polymorphic sites. Pos: genomic position on the reference genome. Ref: reference allele. Alt: alternative allele. Funct. element: functional genomic element. AA residue: amino acid residue position for protein coding genes. aa sub: amino acid residue substitution for protein coding genes. AF: Estimated allele frequency on the set of high quality genomes, expressed as %. FEL: Selection according to FEL: NA=not applicable, yes=under selection, no=not under selection. MEME: Selection according to Meme: NA=not applicable, yes=under selection, no=not under selection. Type: type of selection, NA=not applicable. Completely fixed cluster: list of clusters where the polymorphic site reaches complete fixation (AF>0.9). Fixed supercluster: list of super-clusters where the polymorphic site is fixed (AF >0.01)

Importantly, when the entire collection of 20,521 SARS-CoV-2 genomes considered in this study is taken into account, the total number of variant sites is significantly increased (9398 sites,Supplementary Table S2), but the number and type of high (≥ 1%) frequency variant sites remains relatively constant (46), suggesting a robust estimate of allele frequencies for these sites. Of note, of the 4 sites that show an apparent reduction in prevalence, 3 are located in the 200 nt of the 3’ terminal end of the genome, while one is positioned within the first 36 nt of the 5’ end (Table 1). This suggests that the apparent reduction in allele frequency at these 4 sites is most probably due to an incomplete representation of the terminal ends of the genomes.

Functional annotation of the complete collection of variant sites identified in this study, shows a similar proportion of synonymous and nonsynonymous substitutions among the 6319 low frequency (<1%) variants (50.73% and 49.27% respectively). Conversely, the proportion of non synonymous substitutions is increased to 66.6% (OR=1.31, Fisher p-value 0.159) in the 42 high frequency (≥1%) polymorphic sites associated with protein coding genes (Table 1).

The MEME and FEL methods [47] were applied to the concatenated alignments of protein coding genes of the 11,633 high quality complete genomes to identify signals of adaptive evolution. A total of 194 sites were associated (p-value ≤ 0.05) with signatures of selection according to at least one method (Supplementary Table S3). Of these, 118 sites were associated with positive selection, while 76 sites were deemed to be under negative selection.

Strikingly, a highly significant over-representation of both sites under negative and positive selection is observed among the 42 high frequency polymorphic sites of protein coding genes (Table 1). In particular, 8 of these sites (OR=8.65, Fisher p-value 4.41e-07) are highlighted as evolving under positive selection, while 11 sites (OR=17.82, Fisher p-value 1.95e-12) were predicted to be under negative selection.

### An operational classification of SARS-CoV-2 genomes

An ideal classification system based on molecular data should consider evolutionary features of the species under study to identify the minimum level of divergence required to delineate different types of genomic sequences. Moreover, to avoid excessive-fragmentation each cluster should incorporate a relevant proportion of population. Finally the system should be easy to implement and allow the rapid classification of novel specimens as they become available. Although the classification systems proposed by GISAID and Nextstrain are widely used, we observe that both systems suffer from several limitations, including:

1. **Low reproducibility**: the rules used to establish different clades of genomes are not set out clearly and in a systematic manner. Hence neither existing classification systems can be reproduced/re-applied in full and do not allow the automatic classification of novel strains;
2. **Presence of polyphyletic groups:** although both systems are based on phylogenetic approaches, both the Nextstrain and GISAID proposed classification of SARS-CoV-2 incorporate polyphyletic groups/clusters of genomes (Supplementary Figure S1: see Group G of GISAID; Groups 19A and 20A of Nextstrain);
3. **Bias towards highly abundant genome types:** both in GISAID and Nextstrain groups of genome are established visually based on the relative abundance of groups that populate the phylogeny of SARS-CoV-2. Since sampling of viral genomes is not guaranteed to be uniform across countries, and is indeed biased even between countries with reportedly similar numbers of affected individuals (i.e see for example Italy and UK,Supplementary Table S1) this could result in a over-fragmentation of genome types for which a large number of representative

sequences are available, but also - and more important - to mis-classification of the types of genomes which, for the time being are less represented in the phylogenetic tree. This is exemplified by the fact that at present, more than 1500 genomes are not assigned to a type according to GISAID.

As outlined in the previous section currently available genomic sequences of SARS-CoV-2 show a limited level of variability and a low number of high frequency polymorphic sites. Moreover, a relatively high proportion of these sites may evolve under adaptive selection, possibly consistent with functional consequences of variants. Based on these observations, we propose a simple set of empirical rules for the operational classification of SARS-CoV-2 genomes:

1. Similar to approaches used for Multi Locus Strain typing [42], classification of SARS-CoV-2 genomes should be based on variants that are shared by a relevant proportion of viral population. Since a prevalence of 1% is normally considered to identify genetic variants that have reached fixation in a population [54], we propose to use this cut-off as the natural threshold for identifying high frequency variants.
2. In the light of the fact that closely related genomes show an average of 0.64 polymorphic sites, we suggest that distinct clusters/types should differ at least for 2 high frequency polymorphic sites
3. To avoid an excessive fragmentation, each cluster/type should incorporate at least 100 distinct genomes.
4. Since the variability of SARS-CoV-2 is limited, clusters/types that share one or more high frequency polymorphic sites should form groups of higher order (super-cluster).

To evaluate the validity of the proposed clustering approach, we applied the criteria defined above for clustering both the 11,633 high quality genomes (high quality set), as defined by our stringent criteria, and the entire collection of 20,521 genomes (extended set). Considerations regarding the apparent reduction in allele frequency of polymorphic sites at the ends of the genome (see above), prompted us to exclude these sites from the analyses of the extended set.

Irrespective of the dataset considered (Figure 1A, Supplementary Figure S2A), our criteria consistently identified 12 clusters (C1-C12) and 4 super-clusters (SC1-SC4) (Table 2,Supplementary Table S1). As outlined in Figure 1B (and Supplementary Figure S2B), each cluster and supercluster is defined by a characteristic molecular signature consisting of states at 2 to 9 high frequency variable sites. Cluster 1 is the only exception in this respect, since it is formed by genomes that are highly similar to the reference.

**Table 2:** Comparison of our classification system with GISAID and Nextstrain classification available on June 12th 2020. Size(HQ): number of genomes of the high quality (HQ) dataset of 11,633 genomes assigned to every cluster. Size(complete): number of genomes of the entire dataset (complete) of 20,581 genomes assigned to every cluster. NA= not applicable, the group is not defined.

| **Cluster name** | **Super Cluster** | **Size (HQ)** | **Size (complete)** | **Equivalent group GISAID** | **Equivalent group Nextstrain** |
| --- | --- | --- | --- | --- | --- |
| 1 | 1 | 1140 | 1610 | L | 19A |
| 2 | 2 | 465 | 636 | S | 19B |
| 3 | 2 | 1068 | 1474 | S | 19B |
| 4 | 2 | 124 | 227 | S | 19B |
| 5 | 3 | 3965 | 8102 | GR | 20A+20B |
| 6 | 3 | 133 | 322 | GR | 20A+20B |
| 7 | 3 | 3593 | 5710 | GH | 20A+20C |
| 8 | 3 | 182 | 369 | GH | 20A+20C |
| 9 | 4 | 387 | 928 | V | NA |
| 10 | 4 | 218 | 588 | V | NA |
| 11 | 4 | 154 | 242 | NA | NA |
| 12 | 4 | 204 | 313 | NA | NA |

**Figure 1.**
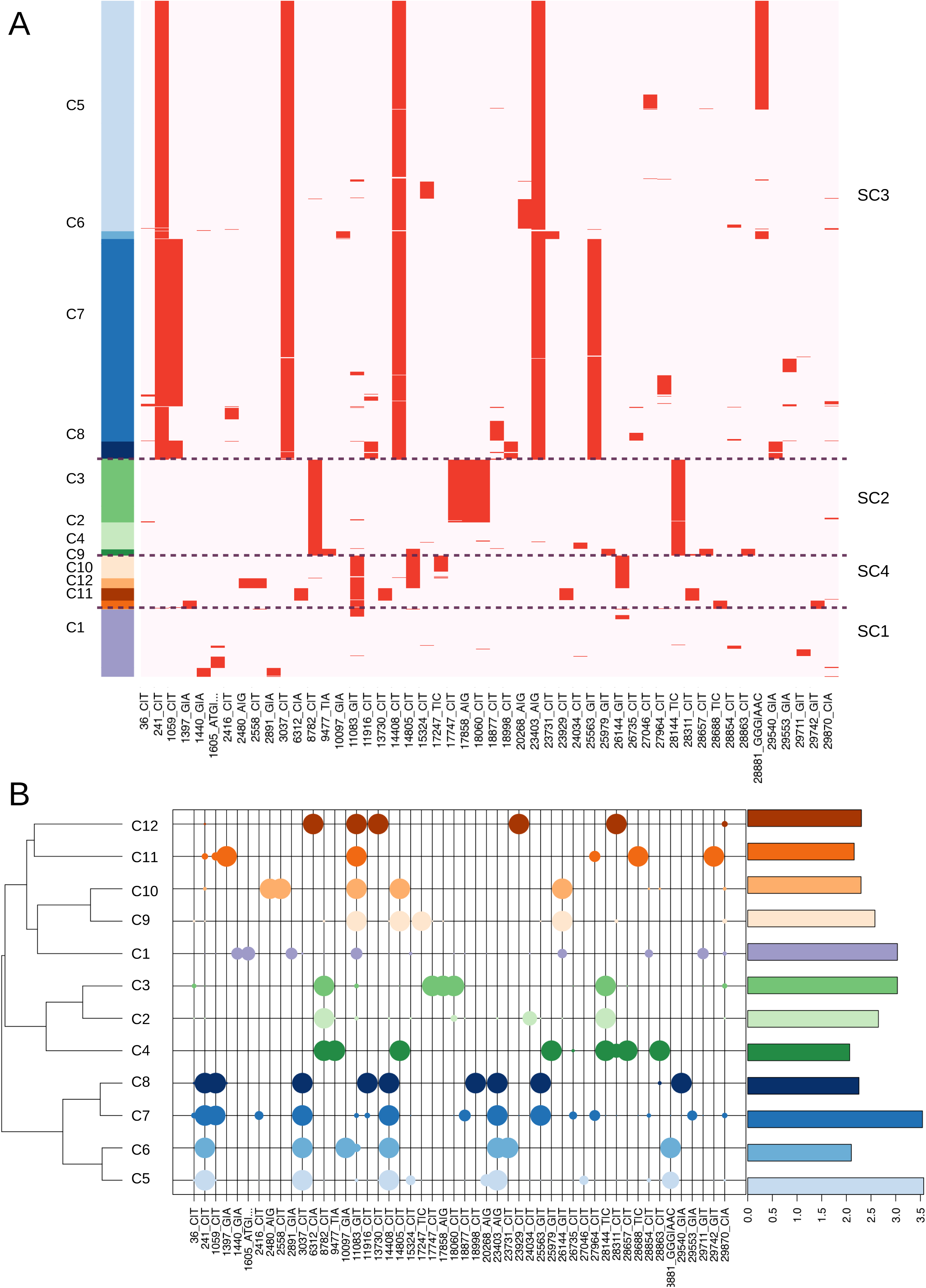
A) Heatmap of presence/absence of 50 high frequency polymorphic sites (AF >0.01) in 11,633 “high quality” complete SARS-CoV-2 genomes, assigned to the 12 clusters here identified. Genomic coordinates are represented on the X axis. Pink indicates a reference allele, Red an alternative allele for that site. The panels on the left indicate cluster memberships, with a different colour assigned to each cluster. Dotted lines delineate super-clusters. B) Bubbleplot of allele frequency of the 50 high frequency polymorphic sites in individual clusters. Color codes according to 1A. The dendrogram on the left indicates clusters with similar allele frequency profiles. The size of each “bubble” is proportional to the frequency of that allele in a given cluster. Barplot on the right panel indicates the number of genomes assigned to every cluster, scaled by logarithm base 10.

A comparison of our proposed clustering systems with the current classifications of Nexstrain and GISAID (Table 2), shows that our method recapitulates all of the major groups of genomes defined by GISAID and Nextstrain, as well as identifies additional clusters undescribed by only one (clusters 9 and 10) or both (clusters 11 and 12) other classification methods. Importantly, although the novel clusters defined in this study are formed by a relatively limited number of genomes (243 and 315 respectively for cluster 11 and 12), they show a similar genetic distance from the reference genome as all the other previously defined larger clusters (Figure 1B). Notably, a heatmap of worldwide prevalence of SARS-CoV-2 types (Figure 2) shows that cluster 12 represents more than 40% of the genomes isolated in India, but shows only a very modest prevalence in all other countries worldwide. To facilitate the classification of newly sequenced SARS-CoV-2 genomes an automated software package that implements the criteria devised in the present study has is made publicly available at https://github.com/matteo14c/assign_CL_SARS-CoV-2.

**Figure 2.**
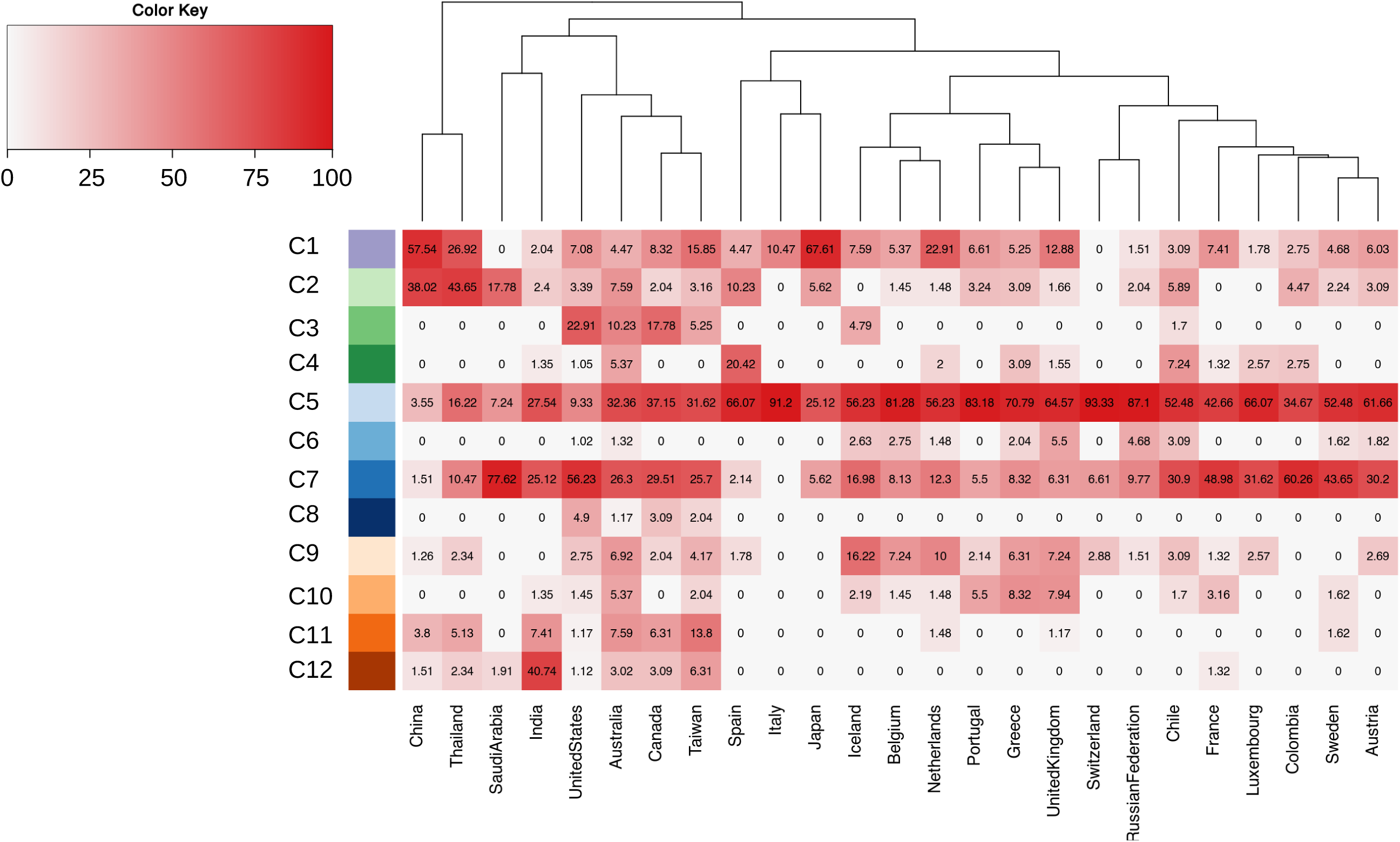
Heatmap of worldwide prevalence of SARS-CoV-2 genome types. Only countries for which at least 50 distinct genomes of SARS-CoV-2 are available in a public repository are shown. Color codes on the left indicate individual clusters, according to Figure 1. The dendrogram on the top delineates groups of countries with a similar prevalence of clusters of SARS-CoV-2 genomes.

Of the 50 high frequency polymorphic sites included in our analyses 35 reach complete, or nearly complete fixation (relative AF ≥ 0.9) in at least one cluster (Table 1, Figure 1B). Of these 35, only three variants (11083G->T; 14805C->T; 28311C->T) show an allele frequency ≥ 0.01 in more than one super-cluster, while the remaining 32 have an AF≥0.01 in only one super-cluster, and can be therefore considered super-cluster “specific”. These observations strongly support our hypothesis that high frequency variable sites, as defined here, are highly effective for the discrimination/classification of distinct types of SARS-CoV-2.

Strikingly, 15 of the 35 sites that are fixed in and specific to at least one cluster are predicted to be under positive (8) or negative (7) selection according to FEL or MEME (Table 1). Although this observation might be suggestive of distinct phenotypic features/properties for the different SARS-CoV-2 types, as previously suggested by other authors [33-35], we remark that in the absence of experimental validations, these data should be interpreted very carefully.

As shown in Figure 3 and consistently with previous reports, we observe a highly biased geographic distribution of SARS-CoV-2 genome types worldwide [27-32]. Indeed, while relatively similar proportions of each cluster/supercluster are observed in Asia, the majority of all viral genomes observed in other continents are assigned to super-cluster 3 (SC3, consisting of C5 to C8). Importantly, from Figure 2 we notice that there is a modest prevalence of SC3 in China, the country that is currently considered the origin of the outbreak, where it accounts for only 5.1 % of all the genome types therein observed.

**Figure 3.**
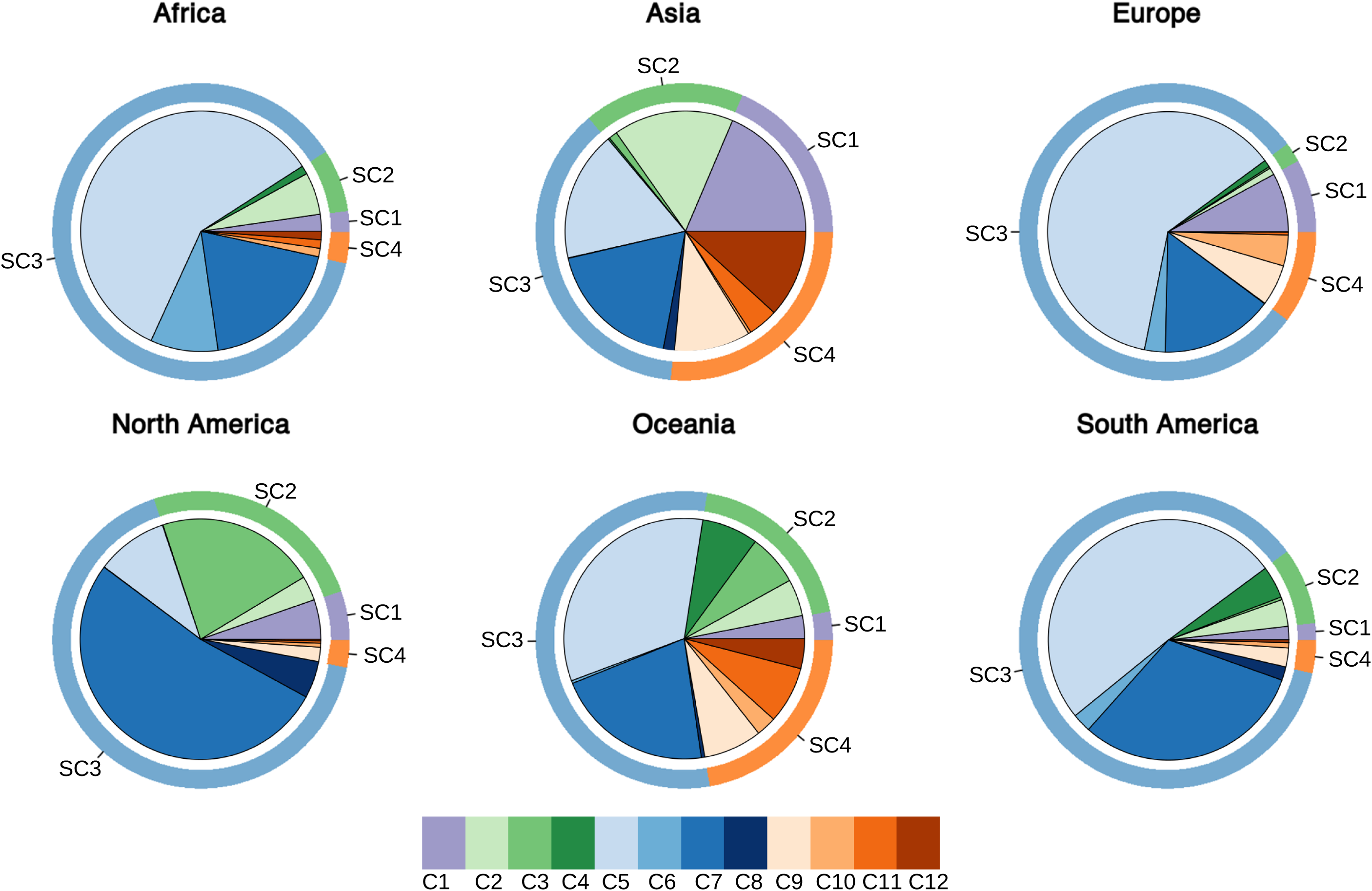
Pie-chart of prevalence of types of SARS-CoV-2 in different continents. Outer circles indicate the prevalence of super-clusters. Color code as in Figure 1

### Spatio-temporal distribution of SARS-CoV-2 genome types and emergence of new types

Phylogeographic analyses show a highly biased distribution of SARS-CoV-2 genomes worldwide (Figure 3). However, representatives of each super-cluster are already observed in different geographic regions of China - the presumed country of origin of the outbreak-within 25 days of the report of the first case of COVID19 in Wuhan (Supplementary Figure S3A). During the same time period, 3 distinct clusters of genomes, each belonging to a different super-cluster (SC1, SC2 and SC4), are already observed in Wuhan (Supplementary Figure S3B). Assuming a constant evolutionary rate of SARS-CoV-2, and based on the average genetic distances of 52 SARS-CoV-2 genomes isolated in China during the early phases of the pandemic (prior to January 20th), we calculated that SARS-Cov-2 might have been circulating undetected in humans at least since August-November 2019, that is 82.3 ± 40.4 days before 26th December 2019, the collection date of the reference SARS-Cov-2 genome (Supplementary Table S4). Strikingly, we notice that, among the 50 high frequency variants of SARS-CoV-2, 12 are also present in one or more genomes of SARSr-CoV-2 isolated from bat and/or pangolin specimens (Supplementary Figure S4). This last observation suggests the presence of a much higher unexplored diversity shared by SARS-CoV-2 and SARSr-CoV-2 viral genomes.

To investigate possible scenarios of emergence of novel genome types, the frequency distribution of the 50 high prevalence alleles (AF ≥ 0.01) was calculated at intervals of 10 days since 26th Dec 2019 (the collection date of the reference genome) separately for each of the 4 super-clusters and for the 12 clusters. Within clusters, distributions of allele frequency are highly stable and do not change over time (Supplementary Figures S5, S6, S7). On the contrary 27 alleles show a rapid emergence and an almost immediate fixation in each viral super-cluster (Figure 4 Supplementary Table S5 Supplementary Table S6). The most notable examples of this pattern are genomic positions 1059, 25563 and 14408 in SC3 and 17747, and 17858 in SC2. Strikingly, 23 out of the 27 alleles rapidly emerging/fixed in super-clusters correspond with polymorphic sites specific to and completely fixed (AF >0.9) within one or more clusters (Supplementary Table S6). This suggests that the early emergence of novel clusters of virus could be used to detect changes of allele prevalence in the whole viral population.

**Figure 4.**
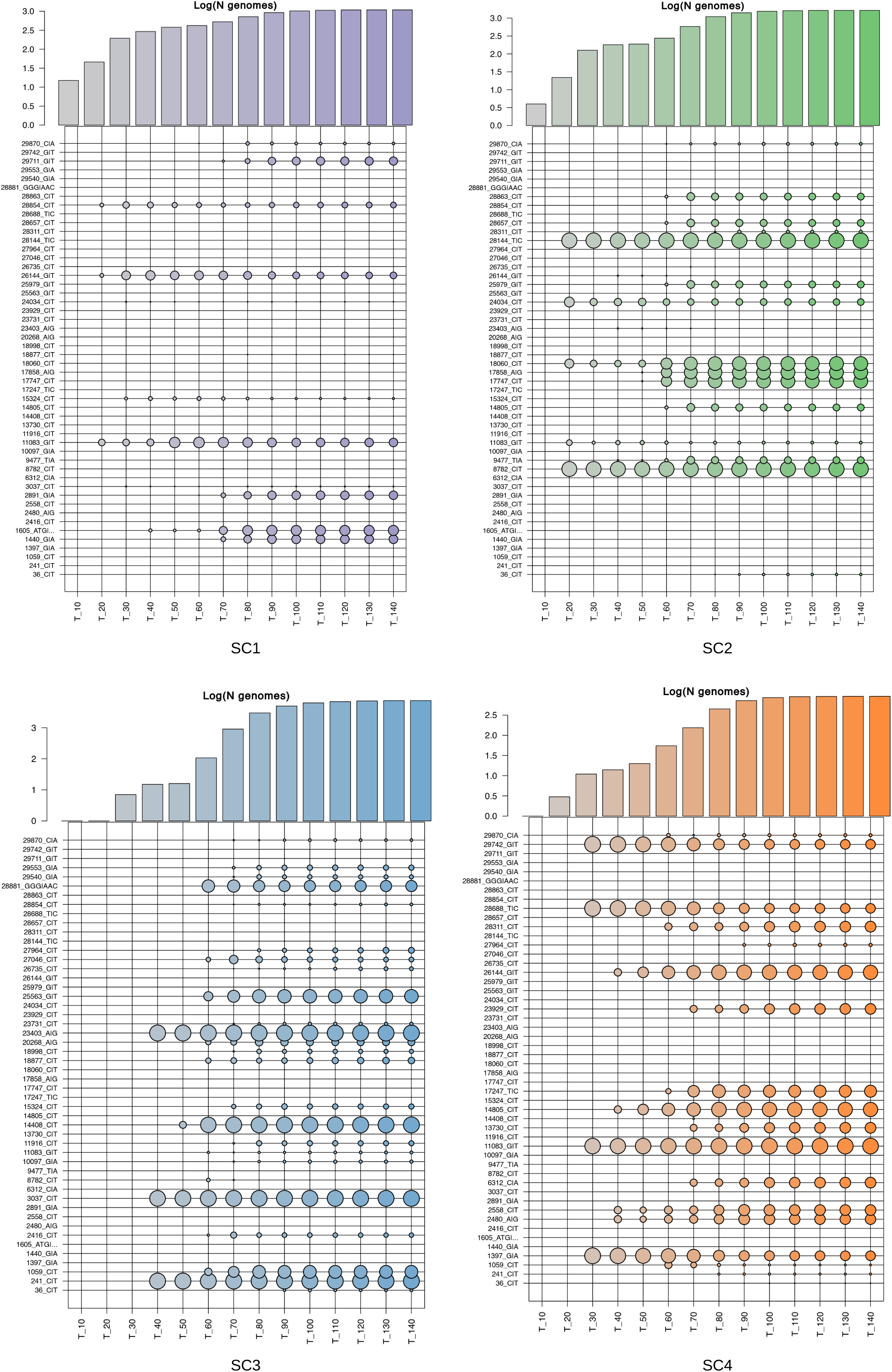
Bubbleplots of allele frequency in the 4 super-clusters of SARS-CoV-2 genomes, at different intervals in time. Each bubbleplot displays the allele frequency of the 50 high frequency polymorphic sites calculated at different-non-overlapped intervals of 10 days. (“T_” with time 0= 26th December 2019). The size of each bubble is proportional to the allele frequency. Color codes according to clusters as in Figure 1. The barplots on the top indicate the number of genomes of each super-cluster observed at each time interval considered, scaled by logarithm base 10.

Strikingly, several clusters of SARS-CoV-2 genomes are not observed during the initial phases of the pandemics and seem to emerge at a later time (Supplementary figures S5, S6, S7). Using an arbitrary threshold of time ≥60 days after the collection of the reference genome (26th Dec 2019), clusters have been divided in “late” (C3, C4, C6, C7, C8, C9, C10, C12: appearing after 60 days) and “early” (C1, C2, C5, C11: appearing before 60 days) (Table 3). We found that “early” clusters have a probable origin in China, as observed in the comparison of the phenetic patterns and the localities of the first 50 isolates (in terms of date) of each cluster (see Supplementary Figure S8). Interestingly, 6 of the 8 “late” clusters are likely to have originally emerged outside China (C4, C6, C7, C8, C10, C12) (Figure 2, Supplementary Figure S9). Overall, the low levels of variability of SARS-CoV-2, the emergence of novel genome types at different times during the pandemic, and the highly biased phylogeographic distribution (Figure 2), suggest that novel genome types of the “late” pandemic phases likely derived from the fixation of novel alleles in “early” pre-existent types.

**Table 3:** Time of emergence and classification of clusters in early and late. Presumed time of emergence: Time of emergence of the cluster, in days with respect to the collection date of the reference genome (time 0 = 26th December 2019). Late/Early: classification of clusters in “early” (emergence of the cluster before time T60), and “late” emergence of the cluster after time T60). Increased variability: P-value for increased variability of evolutionary rates in the cluster according to a Wilcoxon Sum and Rank Test

| **Cluster** | **Presumed time of emergence** | **Late/early** | **Increased variability** |
| --- | --- | --- | --- |
| 1 | 10 | Early | 0.8821 |
| 2 | 20 | Early | 0.8917 |
| 3 | 60 | Late | 0.9132 |
| 4 | 70 | Late | 0.6151 |
| 5 | 40 | Early | 0.667 |
| 6 | 80 | Late | 0.882 |
| 7 | 60 | Late | 0.9146 |
| 8 | 80 | Late | 0.6871 |
| 9 | 60 | Late | 0.8701 |
| 10 | 70 | Late | 0.882 |
| 11 | 30 | Early | 0.8512 |
| 12 | 70 | Late | 0.6912 |

We speculate that several possible alternative evolutionary processes can explain this pattern of rapid allele fixation, including genetic drift and founder effects [55-56], convergent evolution, and/or rapid selection of standing variation for the adaptation to a novel environment [57]. To discriminate between these scenarios, we reasoned that while founder effects and selection should be associated with an overall reduction in genomic diversity of relevant viral sub-populations population, convergent evolution should not alter the underlying allele frequency distribution of the population (except at the sites under selection). We then compared the overall allele diversity between batches of genomes from all the clusters forming the super-clusters SC2 and SC3, batches equivalent both in number and timescale (see Materials and Methods) (Figure 5A). SC1 was not considered due to being composed of a single cluster, while considerations regarding the limited availability of genomes matched by collection date prompted us to exclude SC4 from these analyses. The results show a statistically significant (p-value Wilcoxon 1.3e-07 for C3, 1.5e-07 for C4 and 9.21e-08, 1.21e-07 and 1.13e-07 for C6, C7 and C8 respectively) reduction of genetic diversity for late clusters (i.e. clusters emerging after ≥60 days) compared to early clusters. Importantly (Figure 5B, Table3), we observe that evolutionary rates are highly homogeneous and do not show detectable changes between clusters, suggesting that reduced diversity of late clusters is not associated with a reduction of evolutionary rates. According to our starting hypothesis, and in the light of the strong phylogeographic bias, these results suggest that the emergence of novel SARS-CoV-2 genome types is unlikely to be driven by widespread convergent evolution and independent fixation of advantageous substitutions.

**Figure 5.**
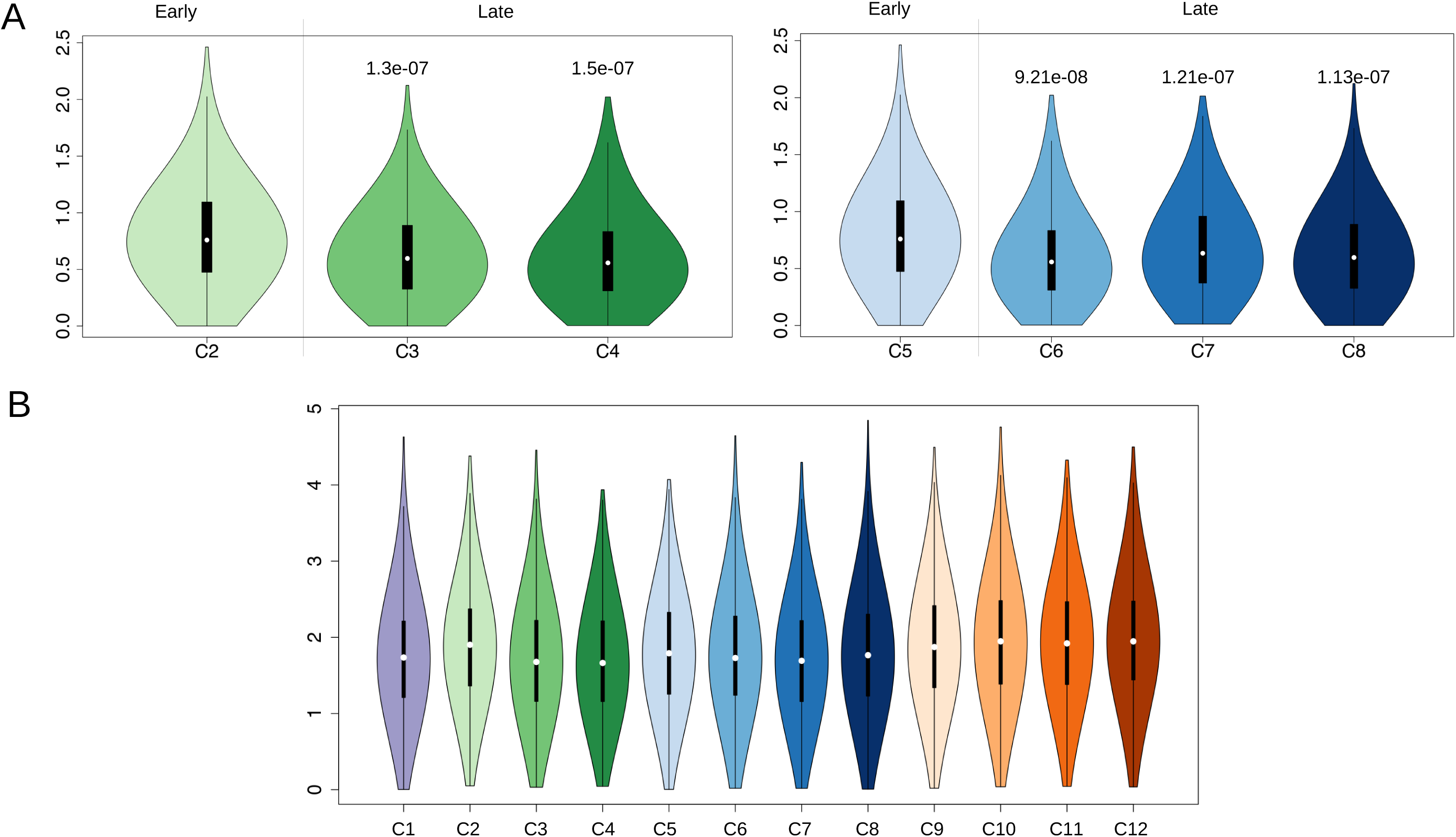
A) Violin plots of genetic diversity of late and early clusters of SARS-CoV-2 genomes assigned to super-cluster CS2 and super-cluster CS3. A vertical line is used to separate early and late clusters. P-values, for the significant reduction of genetic diversity (reduced number of distinct polymorphic sites per genome) are reported on the top of each violin plot. B) Violin plot of substitution rates of the 12 clusters of SARS-CoV-2 genomes identified in this study. Color codes according to Figure 1.

Remarkably, our analyses do not support an increased genomic diversity for clusters included in SC3 (Figure 5A), although their common molecular signature, i.e., the prevalent polymorphic site at position 14408 in the nsp12 gene (RdRp), was previously described as associated with an increased genomic variability (Pachetti et al. [33]). We speculate that biased/incomplete sampling of SC3 during the early phase of the pandemic, and the fact that Pachetti et al compared raw non-normalized genetic distances (instead of normalized evolutionary rates) are the most likely explanation for this discrepancy.

### Distribution of variable sites along the SARS-Cov-2 genome

Patterns of genomic variability were established on sliding windows of 100 bp in size and overlapping by 50 bp, for all the clusters and super-clusters defined in this study. As shown in Figure 6 and Supplementary Figure S10, the observed patterns are remarkably similar between all the clusters and super-clusters, suggesting common patterns of variation. Density of polymorphic sites is significantly enriched (Adjusted Fisher test p-value ≤1e-15 and ≤1e-12 respectively) in both the 5’ and 3’ UTR regions, while protein coding loci (CDS) show less variability. Strikingly, a single genomic region in the 3’ UTR accumulates ∼10x more mutations than any CDS, ∼ 2x more than any other UTR region, and is the single most variable region in the genome of SARS-CoV-2 (Figure 7A). This highly variable genomic region is associated with a conserved secondary structure (Figure 7A), already known in literature and referred to as s2m.

**Figure 6.**
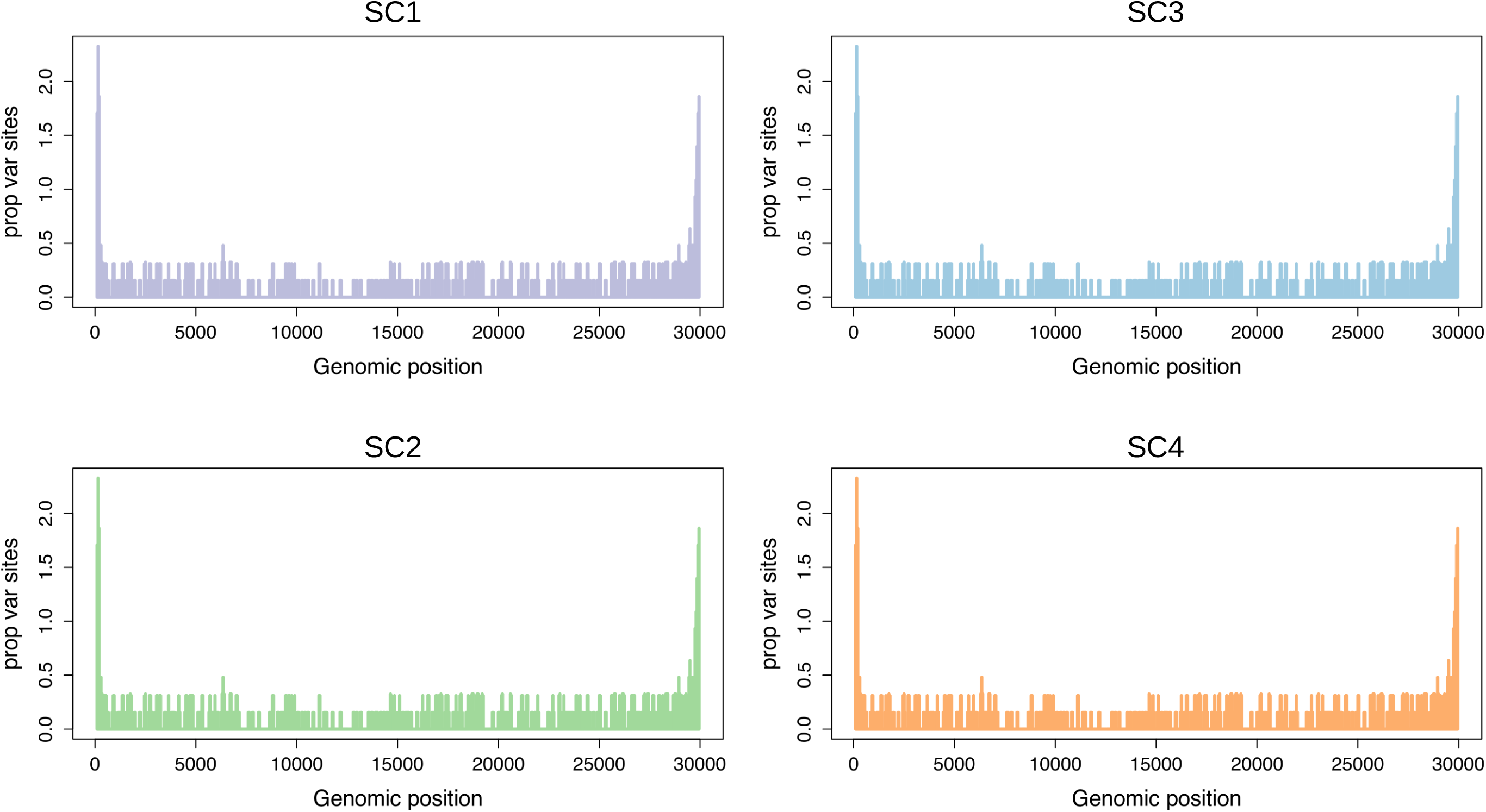
Plot of genomic variability, calculated as the proportion of variable sites identified in overlapping genomic windows of 100 bp in the four super-clusters SC1-SC4. Genomic coordinates are represented on the X axis, number of variable sites per window on the Y axis

**Figure 7.**
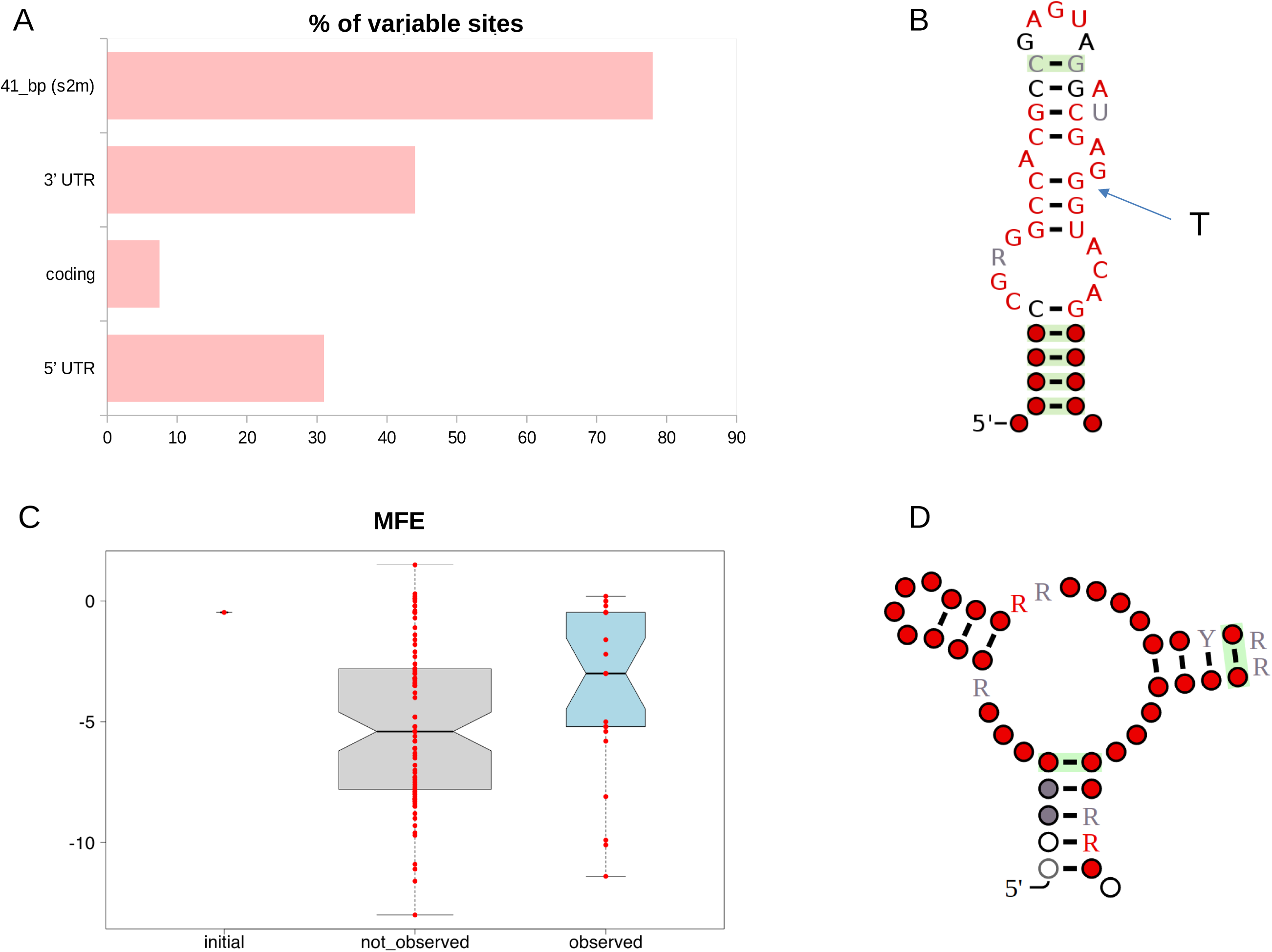
Analysis of variability and structural stability of the s2m secondary structure element. A) Barplot of variability of different categories of genomic elements in the genome of SARS-CoV-2. Variability is reported as the proportion of polymorphic sites. B) Consensus secondary structure of the s2m element of coronaviruses according to the RFAM model RF00164 (https://rfam.org/family/RF00164). The arrow indicates the nucleotide substitution observed in the s2m of the reference genome of SARS-CoV-2 (position 29,758). C) Boxplot of MFE (minimum free energy) of predicted s2m secondary structures. Initial: MFE of the s2m element in the SARS-CoV-2 reference genome. Not observed: MFE of secondary structures associated with single nucleotide substitutions that are not observed in s2m of extant SAR-CoV-2 genomes. Observed: MFE of secondary structures associated with nucleotide substitutions found in the s2m element of extant SAR-CoV-2 genomes D) Prediction of secondary structure co-folding of s2m of SARS-CoV-2 according to the Rscape program. The structure has been derived by alignment of all the 31 distinct variants of the s2m element found in the 11,633 high quality complete genomes considered in this study. Color codes are used to indicate the level of conservation of single nucleotide residues according to the convention used in RFAM (white <=50%, gray >50% and <= 75%, black >75% and < 90%, red>=90%)

S2m is a 41-nucleotide genetic element with a highly conserved secondary - as well as primary and tertiary - structure, and has been described in several families of single-stranded RNA viruses, including Astroviridae, Caliciviridae, Picornaviridae and Coronaviridae [58]. The molecular function of this potentially mobile structural element is not well understood. Current hypotheses include hijacking of host protein synthesis through interactions with ribosomal proteins [59], and RNA interference (RNAi) via processing of the s2m elements into a mature microRNA [60]. In coronaviruses, the highly conserved nature s2m has also allowed the development of a PCR-based virus discovery strategy [61].

As outlined in Figure 7B, compared to the consensus secondary structure of s2m described in the Rfam database, the reference genome of SARS-CoV-2 harbors a nucleotide substitution at a highly conserved and structurally important position, with possible impacts on structural stability (the T at the SARS-CoV-2 genomic position 29,758, indicated by an arrow in Figure 7A). Secondary structure predictions, based on the Vienna RNAfold program, suggest that of all the secondary structures obtained by the possible 129 single nucleotide substitutions in the presumably ancestral sequence of s2m as observed in the genome of RaTG13 SARSr-CoV-2, this would be the second less stable structure in terms of reduction of structural stability (Supplementary Table S7). Based on this observation and on the high levels of variation of the entire s2m region, it is tempting to speculate that s2m could be under diversifying selection in SARS-CoV-2. Strikingly, we observe that the G->T substitution at 29,742, which is a hallmark of cluster C11, would result in a substantially increased stability of s2m (Supplementary Table S7), with an MFE that becomes substantially lower than that of the s2m structure of the reference genome. Conversely, other 2 recurrent substitutions in s2m, 29,742 G->A and 29,734 G->T, which do not reach fixation according to our criteria (having allele frequency ≤ 0.01) and are not associated with a specific cluster of SARS-CoV-2 genomes, are predicted to result only in a marginal decrease of the MFE of the s2m secondary structure (Supplementary Table S7). Interestingly, we notice (Figure 7C) that the same consideration applies to the majority of the nucleotide substitutions that are observed in the SARS-CoV-2 s2m element. Indeed, with respect to the background of all possible nucleotide substitutions that could occur in s2m of the SARS-CoV-2 reference genome, the set of variants that is actually found in extant SARS-CoV-2 genomes seems to produce a less stable secondary structure.

A co-folding analysis of all distinct variants of the s2m elements found in the 11,633 complete and high quality genomes - according to the criteria defined in this study - suggest a very degenerate secondary structure of s2m in SARS-CoV-2 (Figure 7D).

Notably while a substitution that restores the presumably ancestral state of s2m (i.e., the secondary structure of RaTG13 SARSr-CoV-2) is observed (29753 T->G), this substitution is associated only with a very limited number of genomes (AF=0.0012).

## Discussion

Development of effective methods for the study of molecular data and the characterization of pathogens genomic sequences and their evolution are of fundamental importance for monitoring the spread and evolution of pathogens. The GISAID and Nextstrain portals currently represent two invaluable resources for the sharing of genomic data of SARS-CoV-2. Although both GISAID and Nexstrain incorporate methods for the classification of SARS-CoV-2 genomic sequences, these systems are not ideal, since they lack transparent and systematic sets of rules and can not easily be applied for the systematic classification of novel genomic sequences.

In the current study we propose a rational and highly reproducible approach for the classification of SARS-CoV-2 genomes, inspired by Multi Locus Strain Typing (MLST), a molecular approach commonly used for the classification of pathogens [42]. Building on a highly informative set of variable sites, which show high prevalence in the viral population, our system is conceptually simple and represents a general and robust alternative to methods based on phylogenetic analyses. Applying our system to the entire collection of genomic sequences, as available on 28th May 2020, we derive interesting observations concerning evolutionary patterns of SARS-CoV-2.

By comparison of more than 20,000 complete or nearly complete genomic sequences we observe a reduced level of variability and a relatively low mutation rate (1.84 sites per 10^−3^ nt per year) in SARS-CoV-2. Although this is consistent with previous reports of low evolutionary rates in coronaviruses and SARSr-CoVs [26-50], considerations regarding the rapid spread of SARS-CoV-2, and possible biases in sampling of genomes from different areas of the world, suggest that this estimate should be treated carefully, especially since it is not known whether evolutionary rates are expected to remain stable in the event of a rapid expansions of the viral population. This notwithstanding, the presence of different types of SARS-CoV-2 genomes during the early phases of the pandemic (within 25 days of the report of the first case of COVID19 in Wuhan) in several distinct geographic regions of China, coupled with low levels of genomic variability, is suggestive of an early circulation of SARS-CoV-2 in humans, probably well before the major outbreak of COVID19 in Wuhan. The evolutionary rates estimated in this study, coupled with the number of observed polymorphic sites would retrospectively date the spillover of SARS-CoV-2 in humans to August-November 2019, an estimate that is consistent with other recent studies [62]. Careful monitoring of the evolutionary rates of SARS-CoV-2 over a longer period of time, and ideally also on an unbiased/matched number of genomes isolated from different geographic areas, are required to confirm these findings.

In this respect, the fact that a relevant number of SARS-CoV-2 high frequency polymorphic sites are observed even in viral strains isolated from pangolins and bats specimens highlights an unexplored diversity shared by SARS-CoV-2 and SARSr-CoV-2 viral genomes. This suggests that additional sampling of a larger number of non-human specimens is required to reconstruct a more complete phylogeny and to possibly trace back the “original” spillover event. Indeed, notwithstanding the high levels of similarity to SARS-CoV-2 (in the order of 97%), RaTG13, the most closely related viral genome isolated from a bat specimen, is supposed to have diverged from SARS-CoV-2 more than 25 to 40 years ago [63].

Our classification system, based on 50 high frequency polymorphic sites, identifies a total of 12 distinct clusters and 4 super-clusters of SARS-CoV-2 genome types, all having a highly biased phylogeographic distribution (Figure 2-3). We note that several polymorphic sites that are specifically associated (completely fixed) with clusters and super-clusters are predicted to be either under positive or negative selection according to state of the art methods for the study of evolutionary constraints in protein coding genes (Table 1). Interestingly, several of these sites have been previously highlighted by other studies and tentatively associated with increased virulence and/or increased mutation rates of SARS-CoV-2 [33-35].

While fixation of advantageous variants has previously been proposed as an effective and widespread mechanism for the rapid increase of the fitness of a viral population [64], it should be stressed that in the absence of extensive experimental validation, the functional relevance of these genomic variants remains unclear at present. Especially, since reduced levels of variability, high levels of recombination and, particularly, biased sampling of genomic sequences, could impair the accuracy of methods based on phylogenetic reconstruction of ancestral states for the identification of possible evolutionary signatures [65-66]. In this respect, it should be stressed that our observation of reduced genetic variability of “late” viral clusters belonging to SC3 and SC4 (Figure 5A), coupled with the highly biased phylogeographic distribution of SARS-CoV-2 genome types (Figure 2-3), are more consistent with genetic drift and founder effects rather than ongoing adaptive selection. However, the hypothesis that genetic drift accounts for a great part of SARS-CoV-2 variability does not exclude selection having driven a small number of fixed substitutions, so sites identified as candidates for selective evolution warrant further functional characterization both *in vitro* and eventually *in vivo*.

Notably, we observe a highly consistent pattern of nucleotide substitution in SARS-CoV-2 genomes between all clusters and superclusters, characterized by an increased variability at UTRs, in spite of the fact that a significant proportion of genomic assemblies annotated as “full-length” in GISAID are incomplete at the terminal ends. Although this incomplete representation of genomic sequences does not affect the classification system proposed in this study, it might result in an inaccurate/incomplete representation of ongoing evolution of SARS-CoV-2. This is exemplified by the s2m element, a highly conserved secondary structure element located in the 3’ UTR which carries a substitution in the reference genome of SARS-CoV-2 that destabilizes the secondary structure and is possibly deleterious. In the absence of clear data on a function to be experimentally tested, it is very difficult to speculate whether the s2m of SARS-CoV-2 maintains the same functional activity. However, the substantial increase of genomic variability observed in the s2m locus, compared with the rest of the genome, suggests that this element might be subject to ongoing widespread diversifying selection in SARS-CoV-2, or at least have lost significant purifying constraints. Patterns of single nucleotide substitutions in s2m provide contrasting evidence concerning the evolutionary patterns of this secondary structure element in SARS-CoV-2, as the most prevalent substitutions (29,742 G->T) seems to be associated with a considerable increase in secondary structure stability, but the majority of the substitutions observed in extant SARS-CoV-2 genomes are not optimal in terms of the recovery of a highly stable secondary structure.

In conclusion several questions concerning the mechanisms of evolution and the phenotypic characteristics of SARS-CoV-2 (increased/decreased virulence) remain open, in the absence of rigorous experimental validations. However, by allowing a rapid and systematic classification of SARS-CoV-2 genome types, the approach here presented can be extremely useful for a fine grained monitoring of the prevalence of different types of SARS-CoV-2 worldwide, but also for the study of the molecular processes that underlie the emergence of novel viral types

## Acknowledgements

We thank ELIXIR Italy for providing the computing and bioinformatics facilities. We gratefully acknowledge the authors, originating and submitting laboratories of the sequences from GISAID’s EpiFlu™ Database on which this research is based. The list is detailed in Supplementary Table S1

## Supplementary Table Legends

**Supplementary Table S1: Genomic sequence metadata**. High qual: Classification of high quality genomes according to the criteria defined in this work: 1= high quality, 0=not of high quality. Time wrt T0: Time interval in days between the collection date of that genome and 26th December 2019, i.e., the collection date of the reference genome. Release date: date of release in GISAID. Cluster: cluster assigned to each genome according to the classification system defined in this study.

**Supplementary Table S2: Functional annotation of nucleotide variants**. Pos: genomic position on the reference genome. Ref: reference sequence. Alt: alternative sequence. HQ_seq: variant observed in one or more “high quality” genome, 1=yes, 0=no. Number of genomes: Number of distinct genomic sequences that carry the variant. Genomic element: Functional genomic element (protein coding gene or UTR), with numbers corresponding to the relative position within that functional element. AA residue: amino-acidic residue in protein coordinates, NA=not applicable. AA change: Predicted change in protein sequence. NA=not applicable. Predicted FE: Predicted functional effect, with FDdel= Frameshift deletion, FSins=Frameshift insertion, NS=non synonymous substitution, S=synonymous substitution, Stop Gain= stop gain, Stop loss= stop loss. MEME/FEL: evidence for selection according to MEME or FEL.

**Supplementary Table S3: List of amino acid residues under selection according to MEME and FEL.** FEL: Under selection according to FEL 1=True, 0=False. MEME: Under selection according to MEME 1=True, 0=False. Kind: Type of selection (Positive/negative).

**Supplementary Table S4: Average genetic distances and estimated divergence times of 52 genomic sequences isolated in China within 25 days of the collection of the reference genome.** AvG dist: Average genetic distance. Avg dist months: Average distance in months. Interval: Interval (in days) between the collection date of that genome and 26th December 2019, i.e., the collection date of the reference genome.

**Supplementary Table S5: Allele frequency of the 50 high frequency polymorphic sites in the 4 super-clusters of genomes defined in this study.** “T_” indicates time expressed in days, with time 0= 26th December 2019, i.e., the collection date of the reference genome

**Supplementary Table S6: List of 26 genetic variants fixed in a super-cluster after 60 or more days from the collection date of the reference genome (26th December 2019).** Time: Time of fixation of the variant. AV-Prev: Average allele frequency in the super-cluster. VAR-Prev: Variance of allele frequency in the super-cluster. Completely fixed: Complete fixation in one or more clusters (AF>0.9),1=True, 0=False

**Supplementary Table S7: Analysis of stability of secondary structure of the s2m element.** Rel pos: relative position in s2m. Gen pos: genomic position. Sub: Nucleotide substitution. MFE: MFE. Observed: Observed in an actual genome sequence: 1=True, 0=False. Sub_RaTG13: Nucleotide substitution in the s2m of the RaTG13 genome. MFE_RaTG13: MFE in the s2m of the RaTG13 genome. The first row indicates the MFE (minimum free energy) of s2m secondary structure found in the reference genome of SARS-CoV-2 and in the RaTG13 SARSr-CoV-2 genome assembly. NA=not applicable

## Supplementary Figure legends

**Supplementary Figure S1:** Phylogeny and classification of SARS-CoV-2 genomes according to the GISAID and Nextstrain portals. A) Nextstrain, https://nextstrain.org/ncov/global accessed June 12th 2020 B) GISAID https://www.epicov.org/epi3/frontend#, from GISAID “analysis update”, accessed June 12th 2020.

**Supplementary Figure S2:** A) Heatmap of presence/absence of 46 high frequency polymorphic sites (AF >0.01) in the complete collection of 20,521 SARS-CoV-2 genomes, assigned to the 12 clusters here identified. Genomic coordinates are represented on the X axis. Pink indicates a reference allele, Red an alternative allele for that site. The panels on the left indicate cluster memberships, with a different colour assigned to each cluster. Dotted lines delineate superclusters.

B) Bubbleplot of allele frequency of the 46 high frequency polymorphic sites in individual clusters. Color codes according to Figure 1A. The dendrogram on the left indicates clusters with similar allele frequency profiles. The size of each “bubble” is proportional to the frequency of that allele in a given cluster. Barplot on the panel indicates the number of genomes assigned to every cluster, scaled by logarithm base 10.

**Supplementary Figure S3:**

A) Prevalence of super-clusters and clusters in four distinct regions of China, for which more than 20 genomic sequences were collected within 25 days of the collection date of the reference genome (26th December 2019). Pie-charts represent prevalence of clusters. Outer circles indicate the prevalence of super-clusters.

B) Prevalence of super-clusters and clusters in Wuhan city within 25 days of the collection date of the reference genome (26th December 2019). Pie chart as described above. In the heatmap genomic coordinates are represented on the X axis. Pink indicates a reference allele, red an alternative allele for. Panels on the left indicate cluster memberships, with a different colour assigned to each cluster. Color codes according to Figure 1. Collection dates are reported on the rows.

**Supplementary Figure S4: Phenetic patterns of the 50 high frequency SARS-CoV-2 alleles, in SARSr-CoVs-2**. SARS-CoV-2 genomic positions for the 50 high frequency (AF>=0.01) polymorphic sites of SARS-CoV-2 are represented on the X axis. Pink indicates a reference allele, red an alternative allele. GISAID accession numbers and species (pangolin or bat) of SARSr-CoVs-2 sharing some of these sites are reported on the rows.

**Supplementary Figure S5: Bubbleleplot of allele frequency in individual clusters of super-cluster SC2.** Allele frequency of the 50 high frequency polymorphic sites in super-cluster SC2, calculated at different and non-overlapped intervals of 10 days (“T_”, with time 0 = 26th December 2019, i.e., the collection date of the reference genome). Color codes according to Figure 1. The barplots on the top indicate the number of genomes of each super-cluster observed at each time interval considered.

**Supplementary Figure S6: Bubbleplot of allele frequency in individual clusters of super-cluster SC3.** Allele frequency of the 50 high frequency polymorphic sites in super-cluster SC3, calculated at different and non-overlapped intervals of 10 days (“T_”, with time 0 = 26th December 2019, i.e., the collection date of the reference genome). Color codes according to Figure 1. The barplots on the top indicate the number of genomes of each super-cluster observed at each time interval considered.

**Supplementary Figure S7: Buppleplot of allele frequency in individual clusters of super-cluster SC4.** Allele frequency of the 50 high frequency polymorphic sites in super-cluster SC4, calculated at different and non-overlapped intervals of 10 days (“T_”, with time 0 = 26th December 2019, i.e., the collection date of the reference genome). Color codes according to Figure 1. The barplots on the top indicate the number of genomes of each super-cluster observed at each time interval considered.

**Supplementary Figure S8: Phenetic patterns and geographic area of the first 50 genomes for each “early” cluster.** Each heatmap displays presence/absence of the 50 high frequency polymorphic sites (AF >0.01, identified in the 11,633 “high quality” complete SARS-CoV-2 genomes) in the first 50 genomes, in terms of date, of each early cluster. Genomic coordinates are represented on the X axis. Pink indicates a reference allele, red an alternative allele for that site. Early cluster as defined in Table 3. Panels on the left indicate continents: yellow=Asia, blue=Europe, pink=Africa, green=North America, orange=South America, purple=Oceania. Country and date of collection are reported on the rows.

**Supplementary Figure S9: Phenetic patterns and geographic area of the first 50 genomes for each “late” cluster.** Each heatmap displays presence/absence of the 50 high frequency polymorphic sites (AF >0.01, identified in 11,633 “high quality” complete SARS-CoV-2 genomes) in the first 50 genomes, in terms of date, of each late cluster. Genomic coordinates are represented on the X axis. Pink indicates a reference allele, red an alternative allele for that site. Panels on the left indicate continents: yellow=Asia, blue=Europe, pink=Africa, green=North America, orange=South America, purple=Oceania. Country and date of collection are reported on the rows.

**Supplementary Figure S10:** Plot of genomic variability, calculated as the proportion of variable sites identified in overlapping genomic windows of 100 bp in the twelve clusters C1-C12. Genomic coordinates are represented on the X axis, number of variable sites per window on the Y axis. Color codes according to Figure 1.

