## Supplementary figures and images for "Comparative genomics provides an operational classification system and reveals early emergence and biased spatio-temporal distribution of SARS-CoV-2"

### Supplementary Figure S1

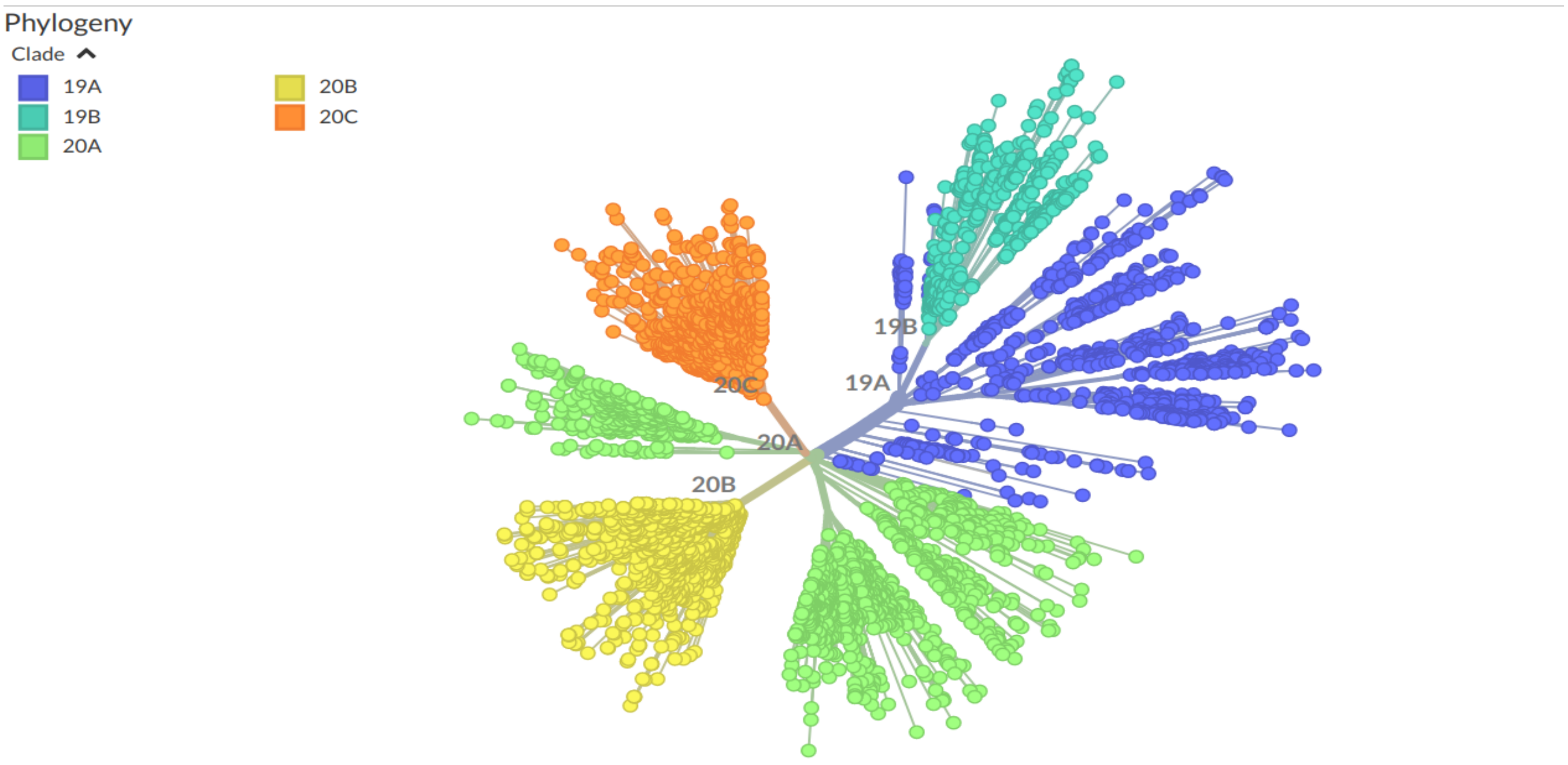

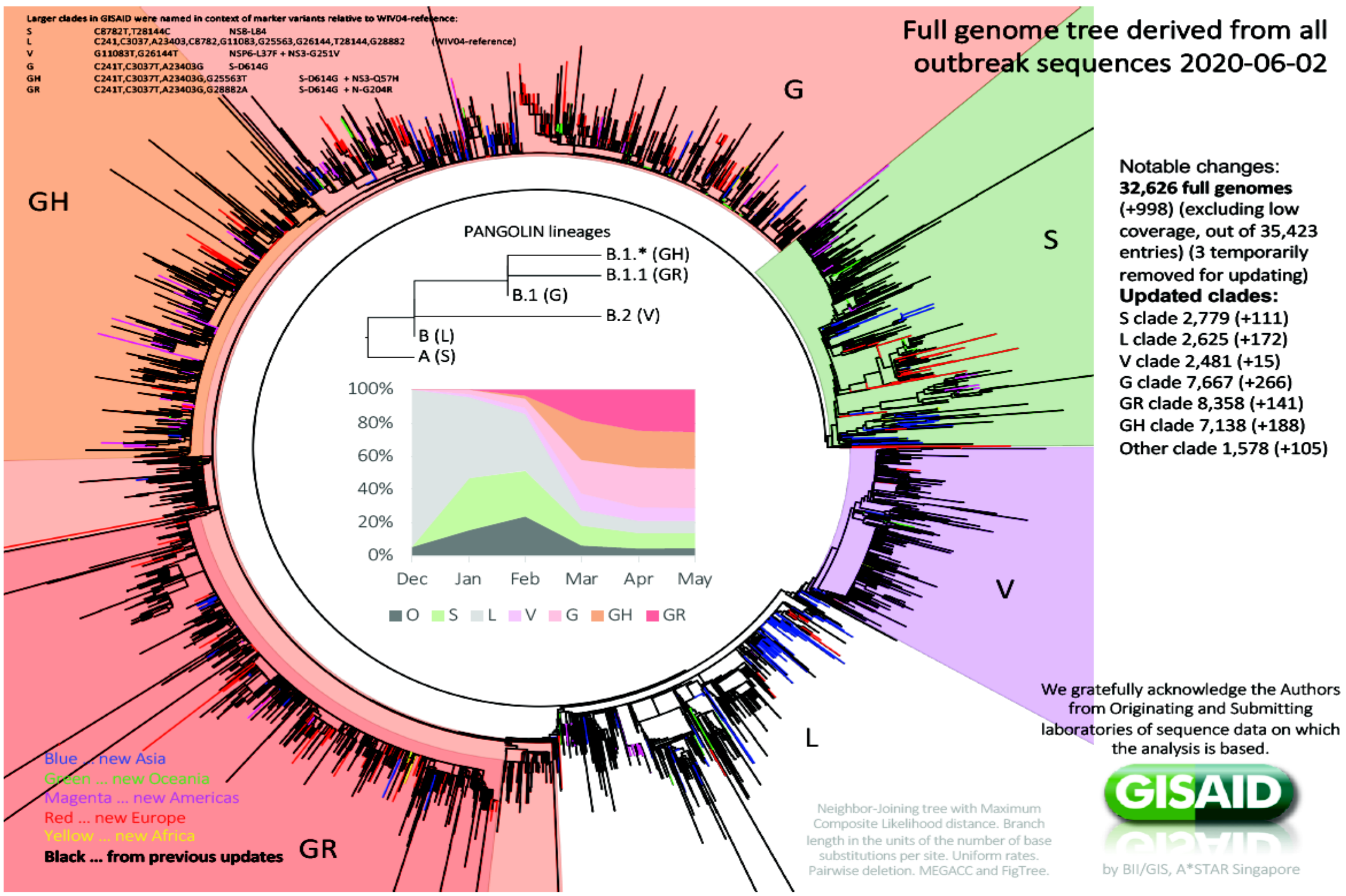

### Supplementary Figure S2

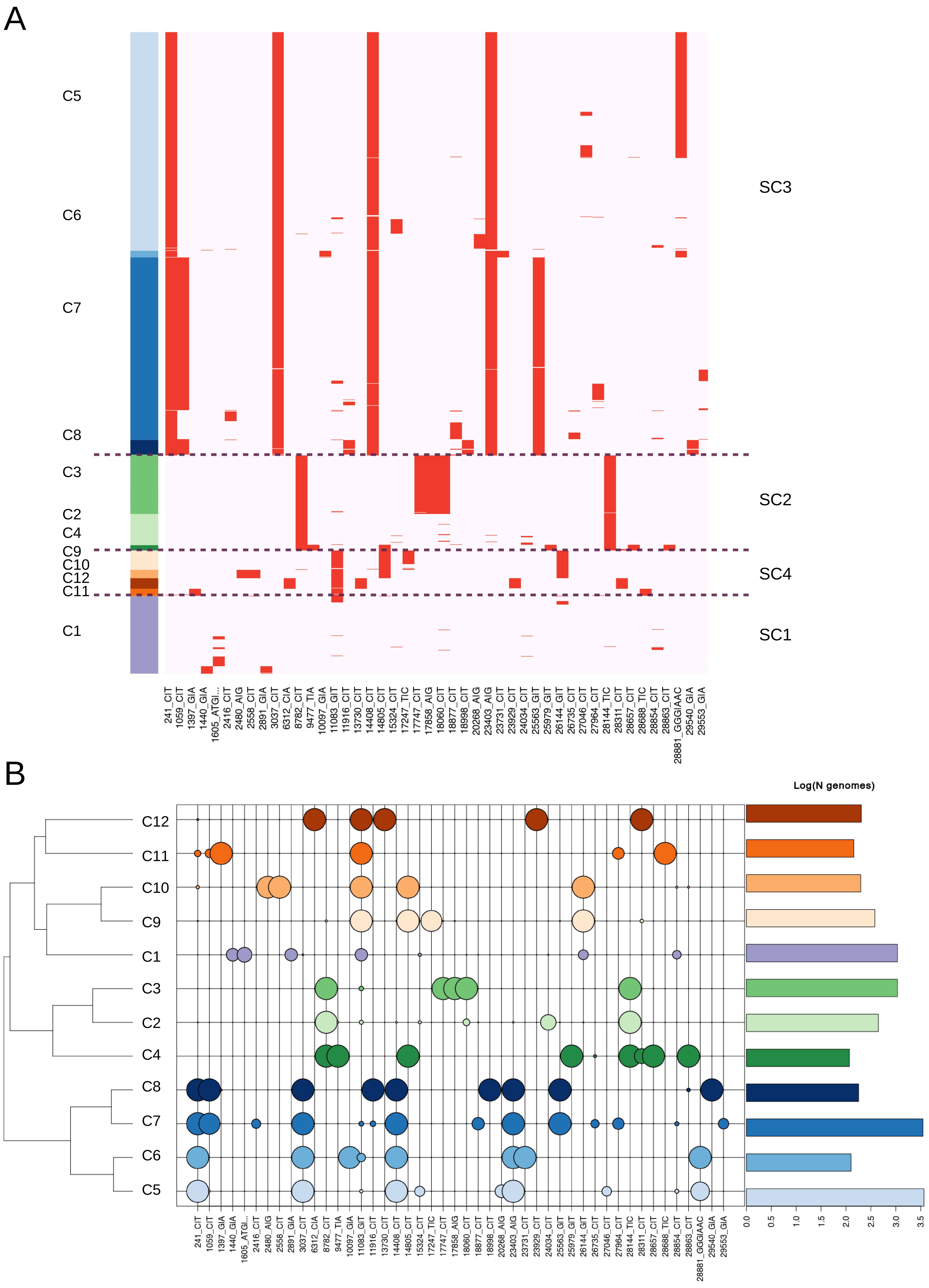

### Supplementary Figure S3

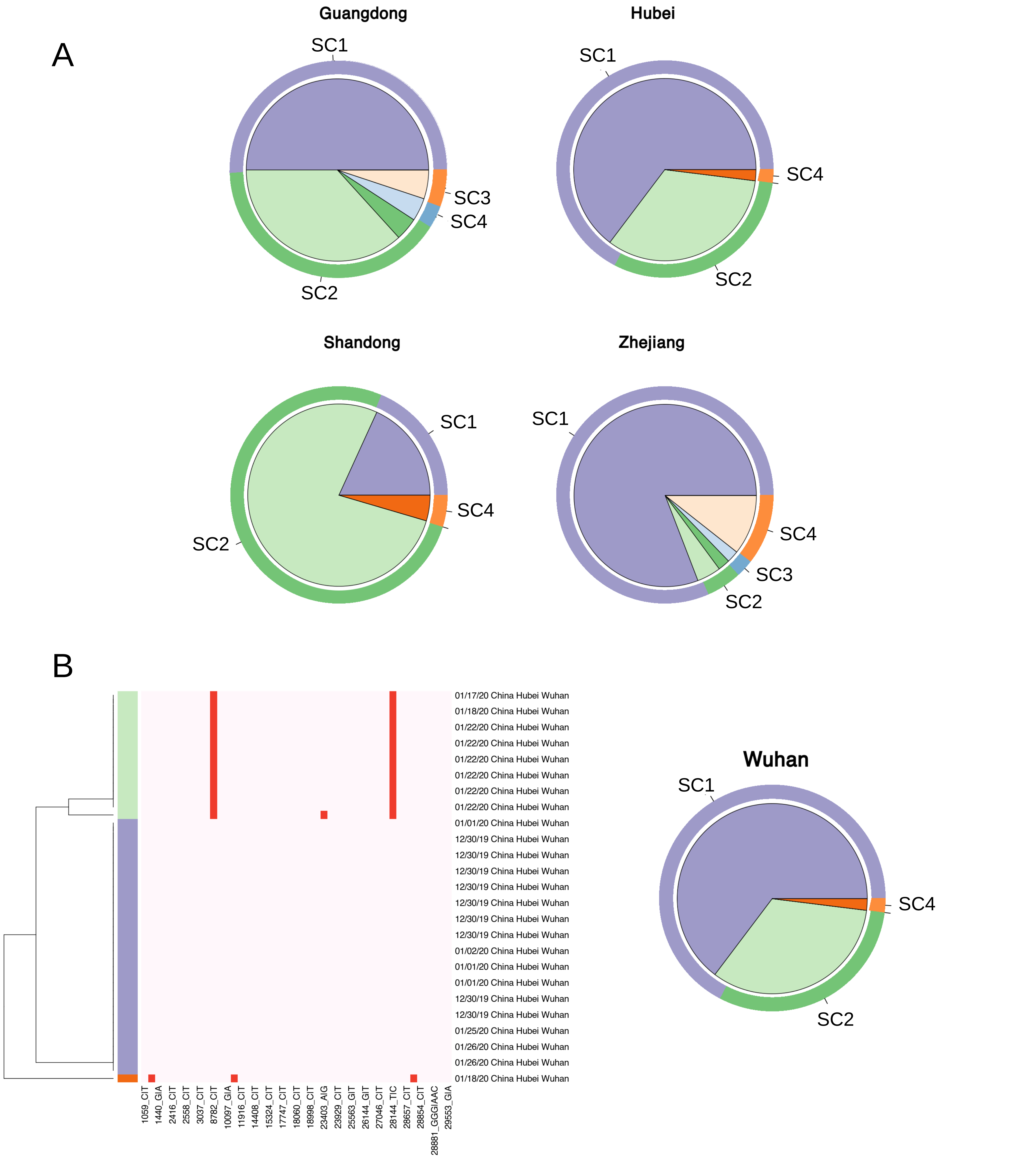

### Supplementary Figure S4

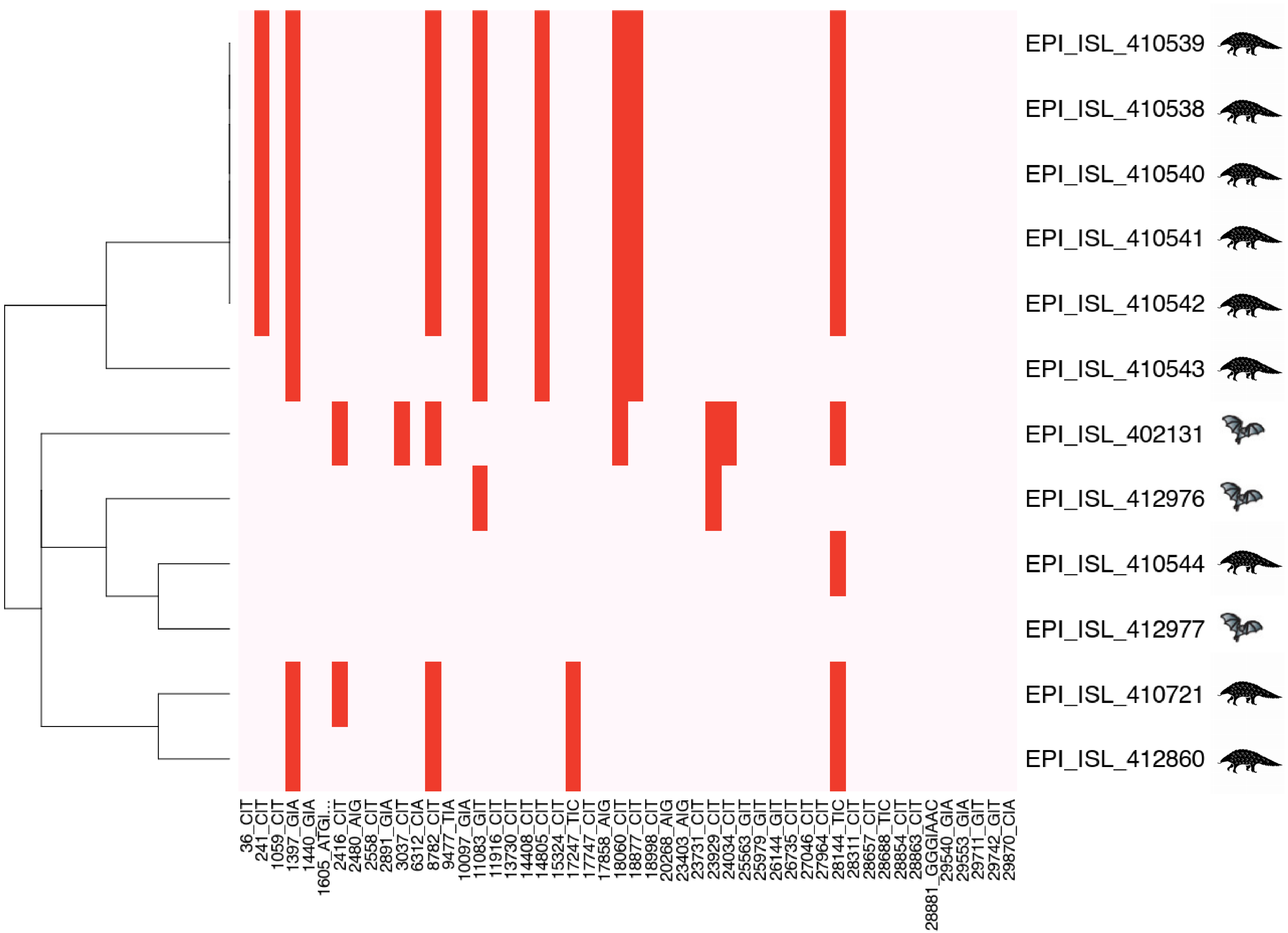

### Supplementary Figure S5

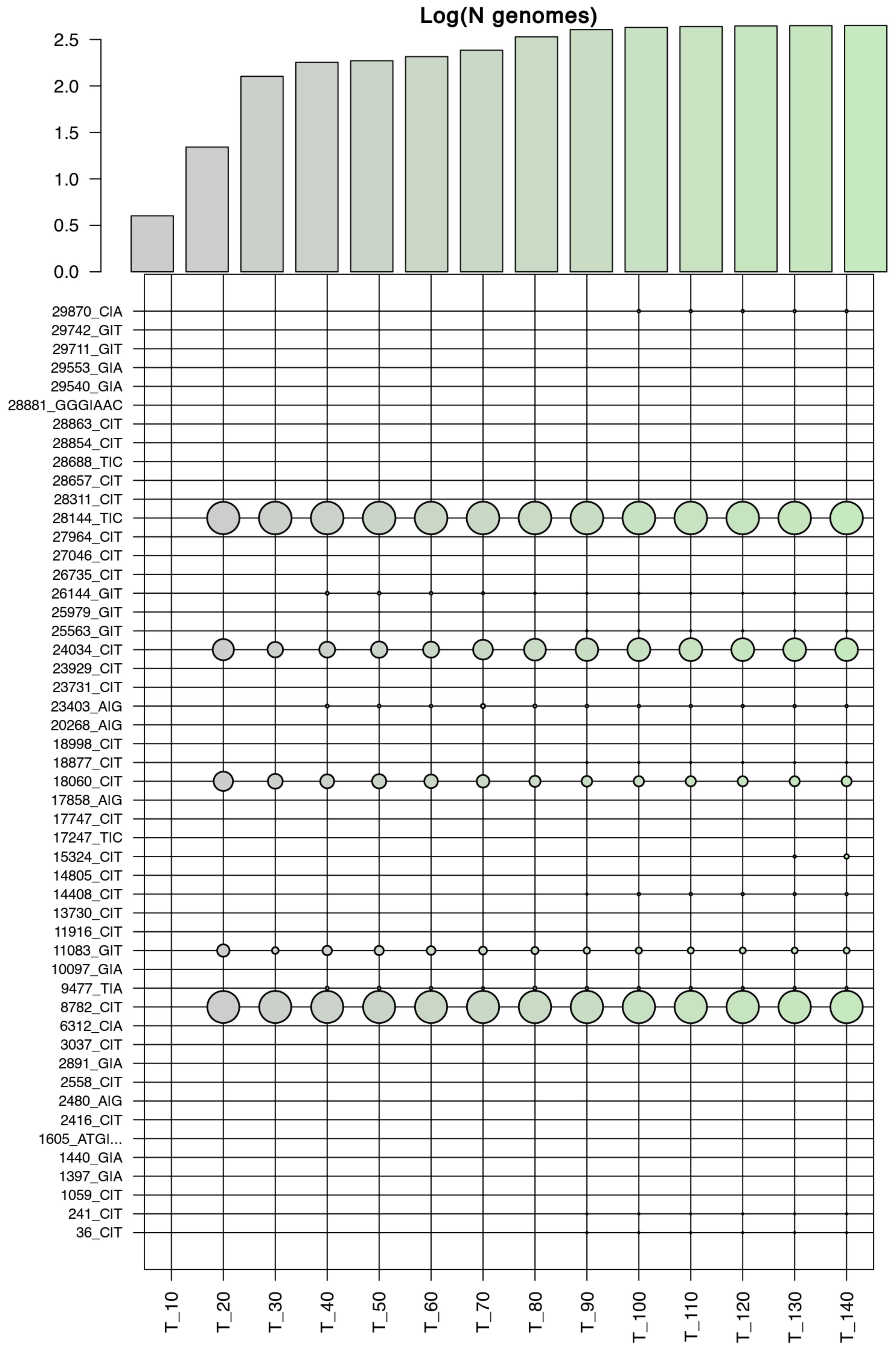

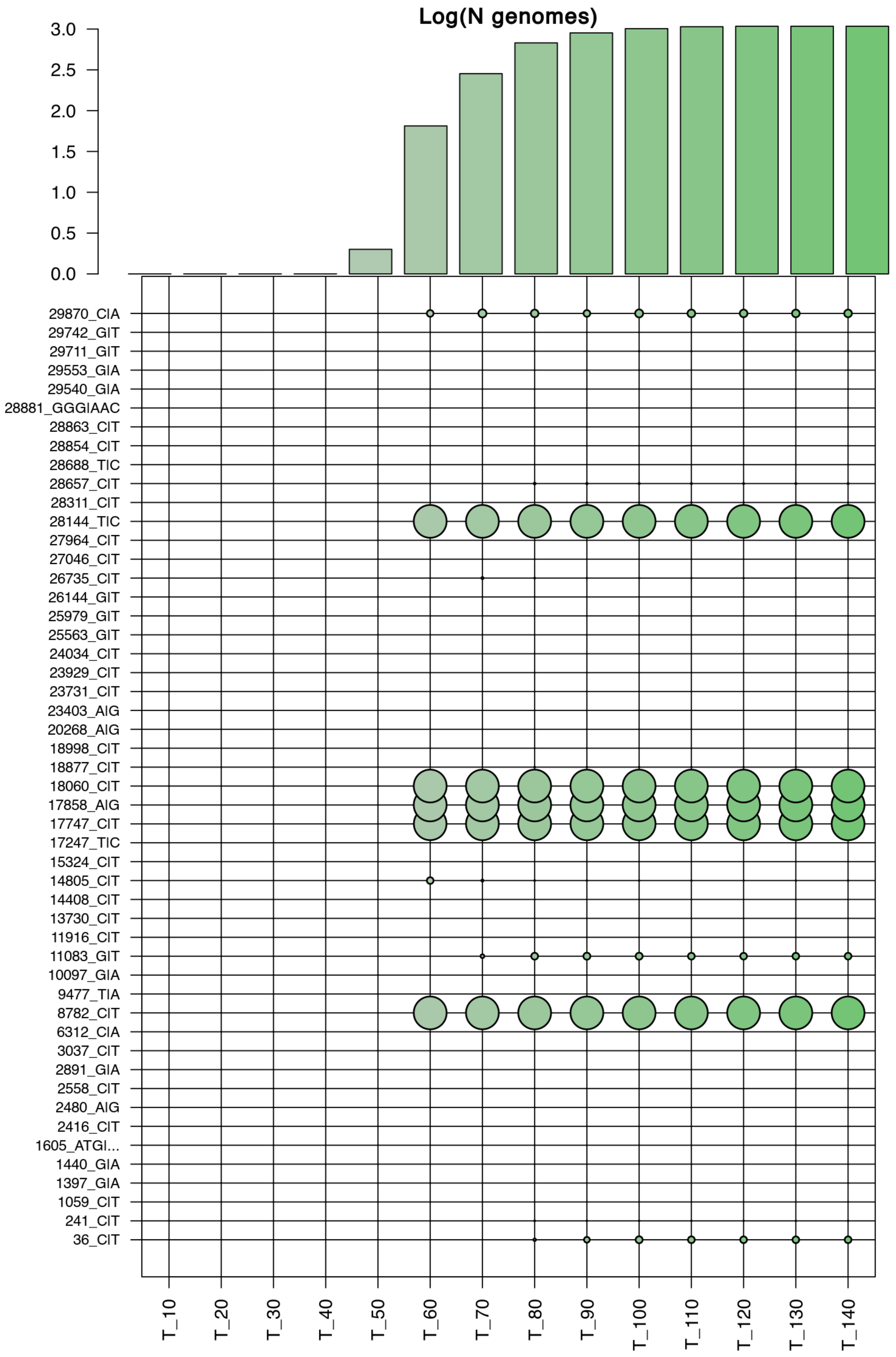

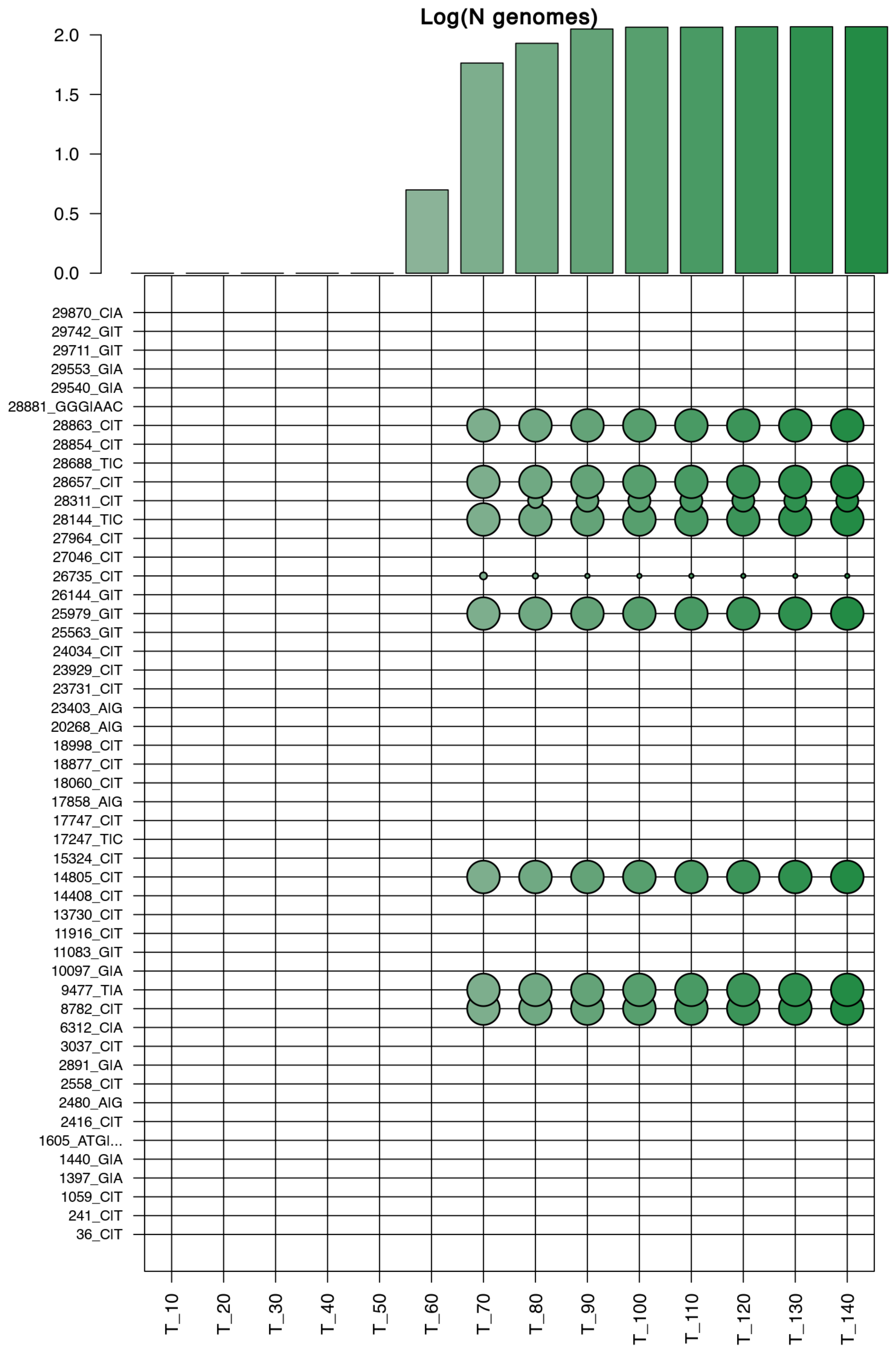

### Supplementary Figure S6

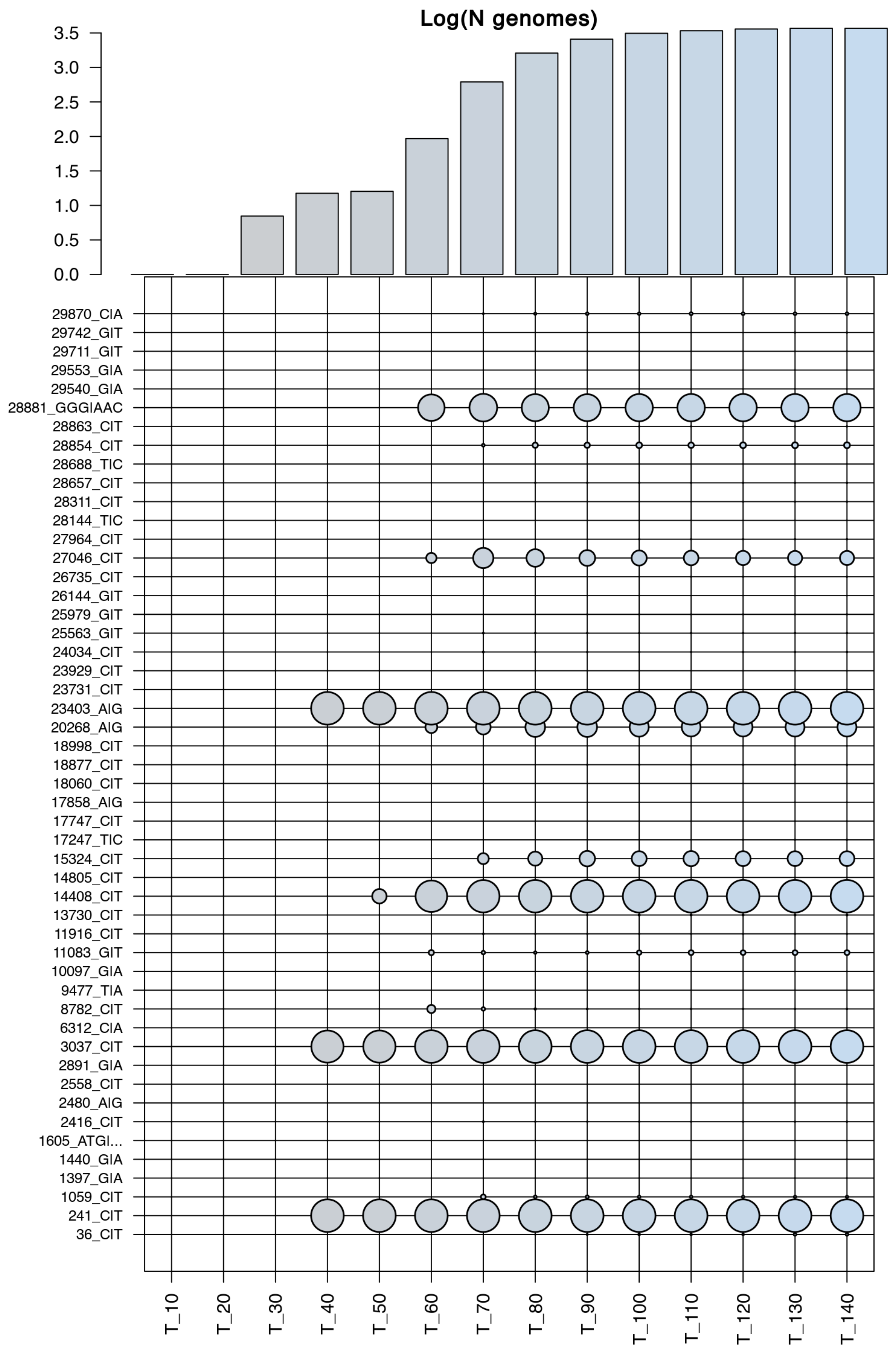

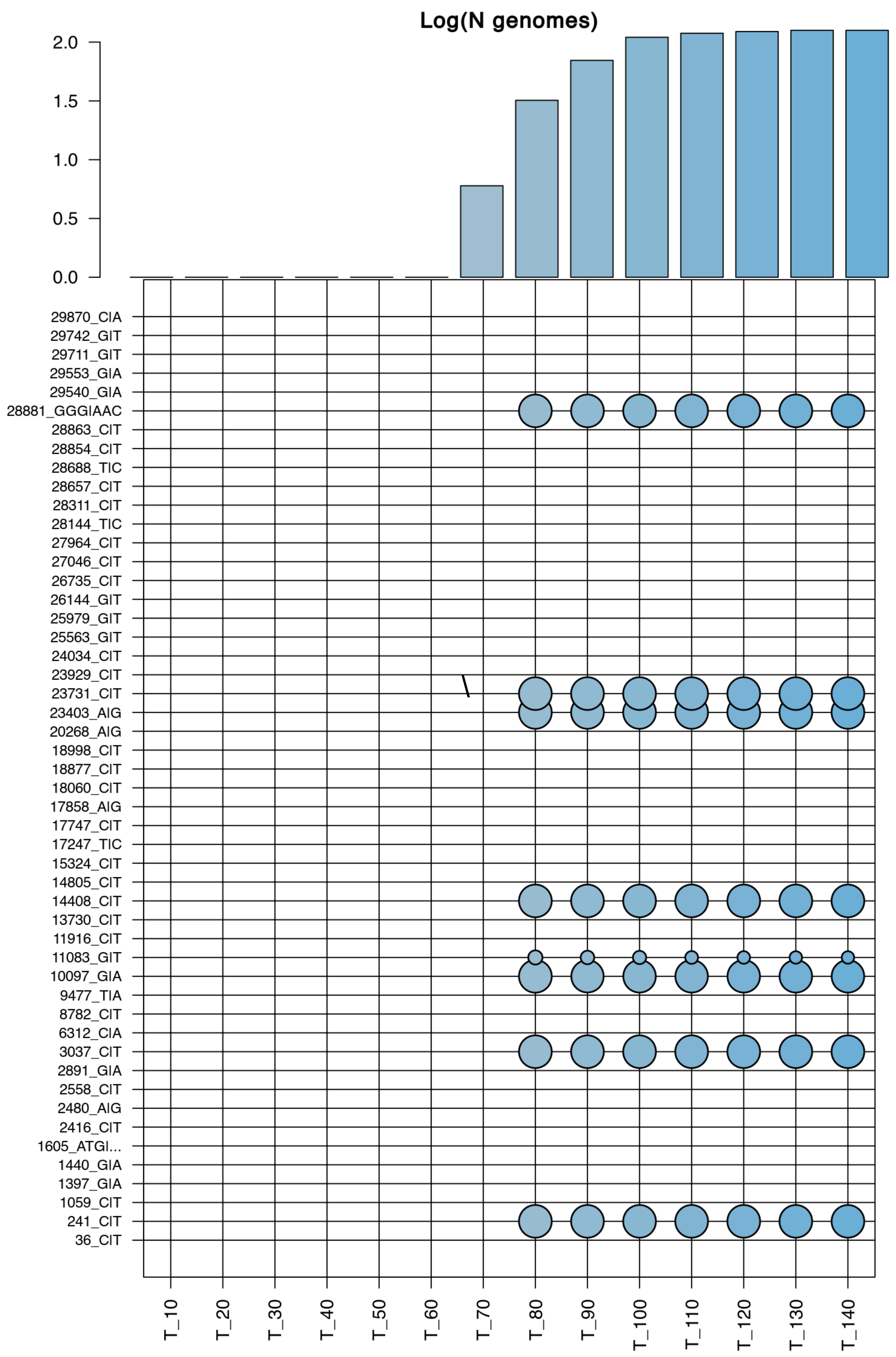

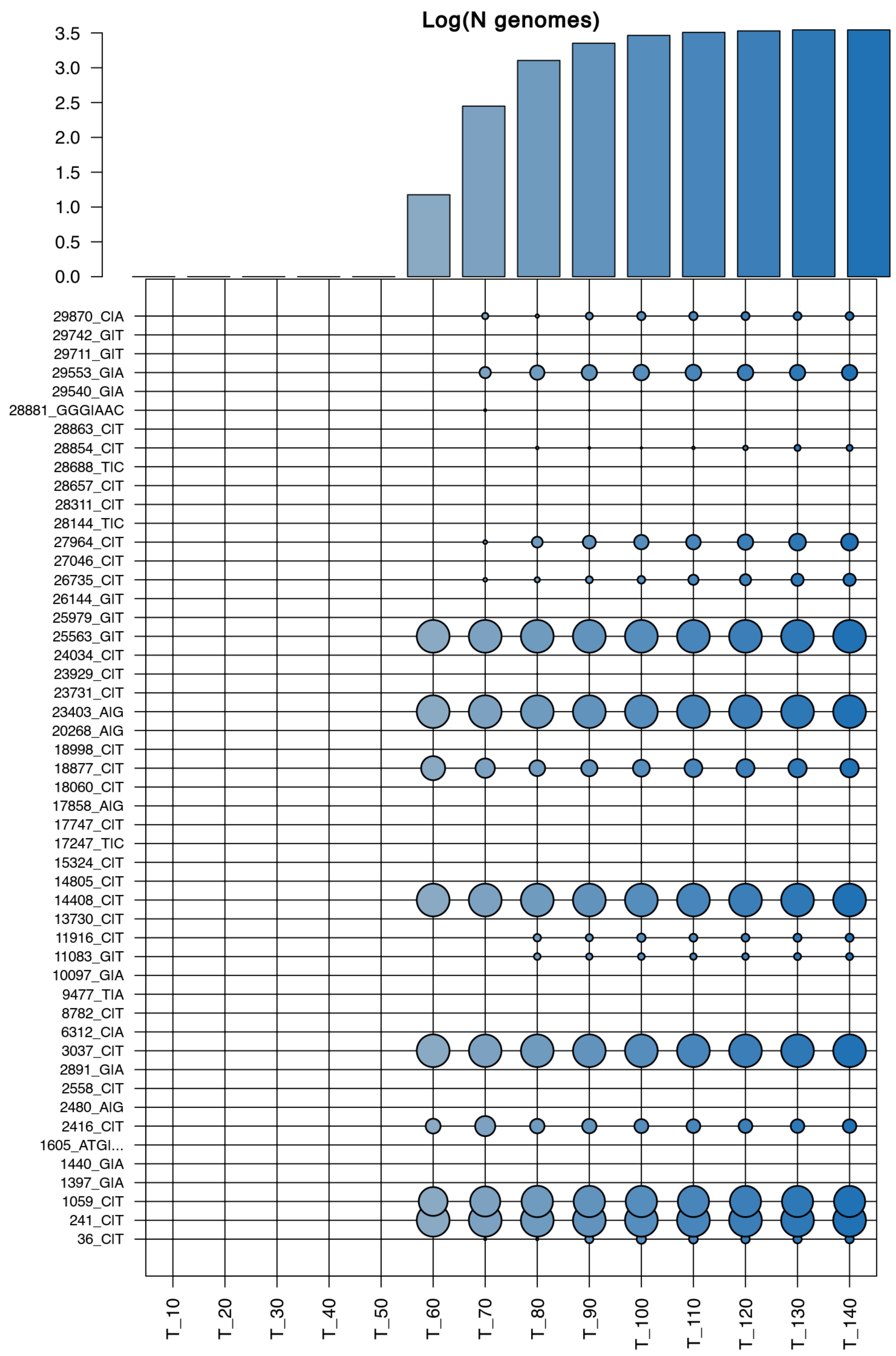

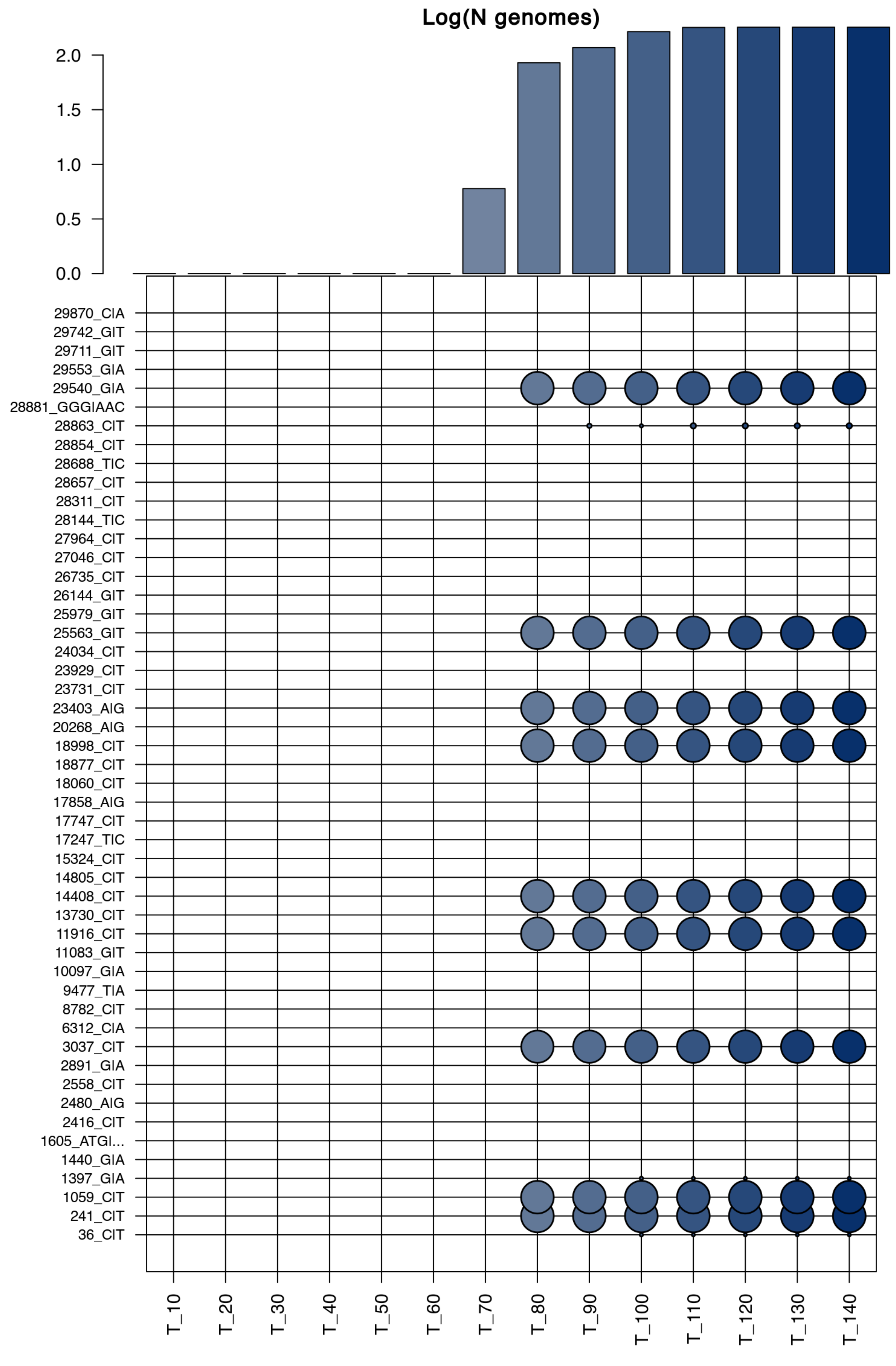

### Supplementary Figure S7

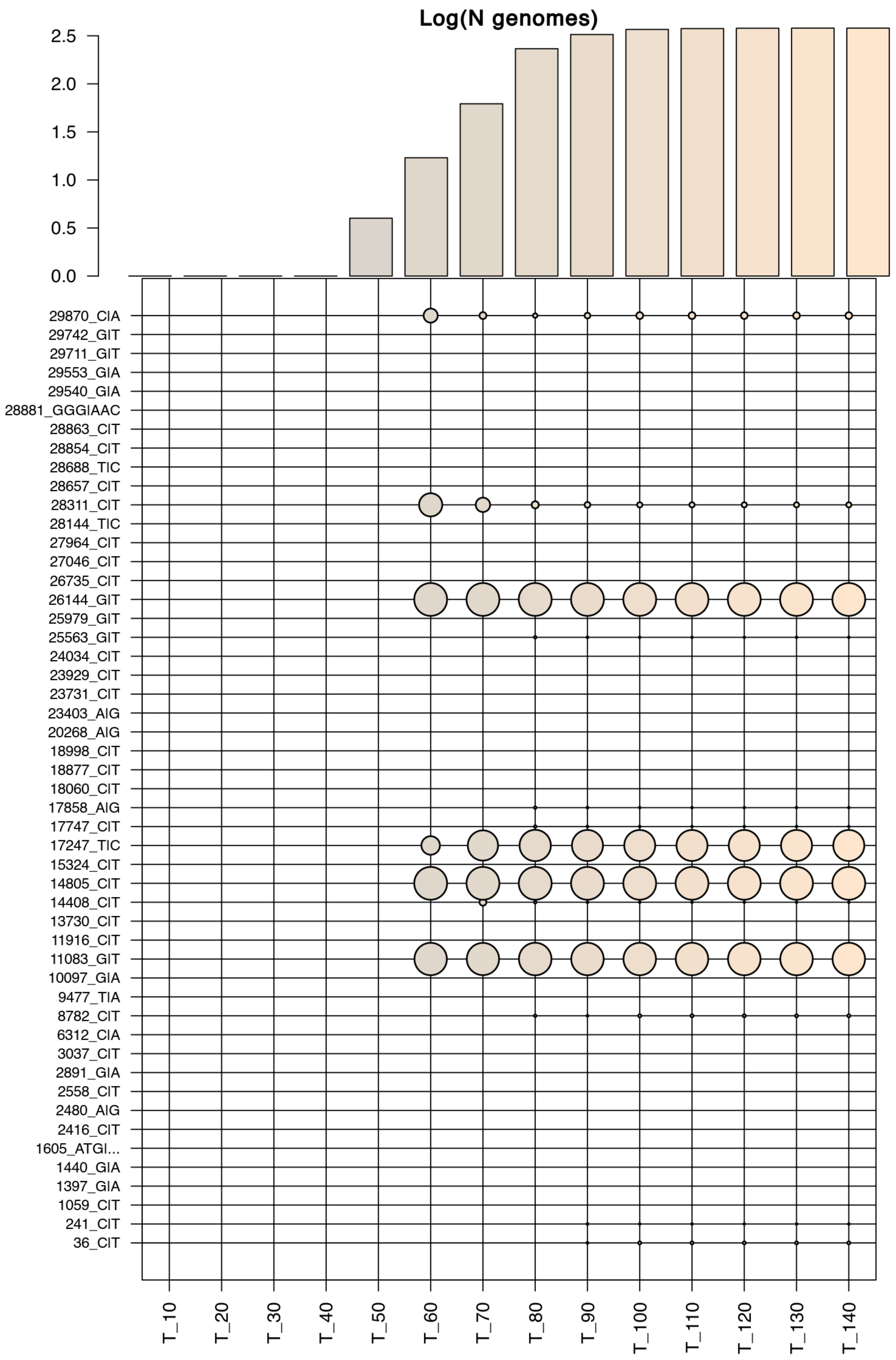


C9

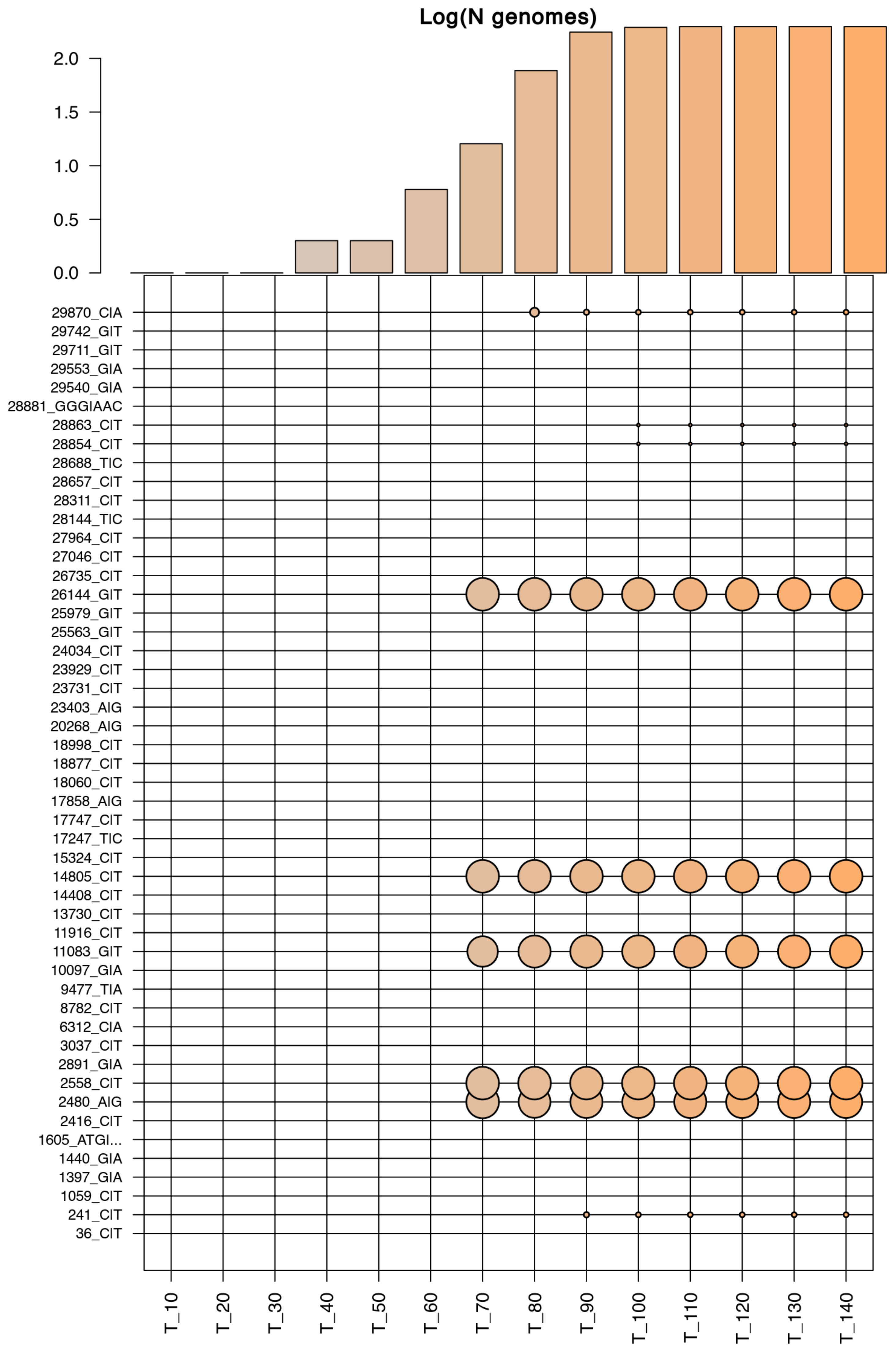


C10

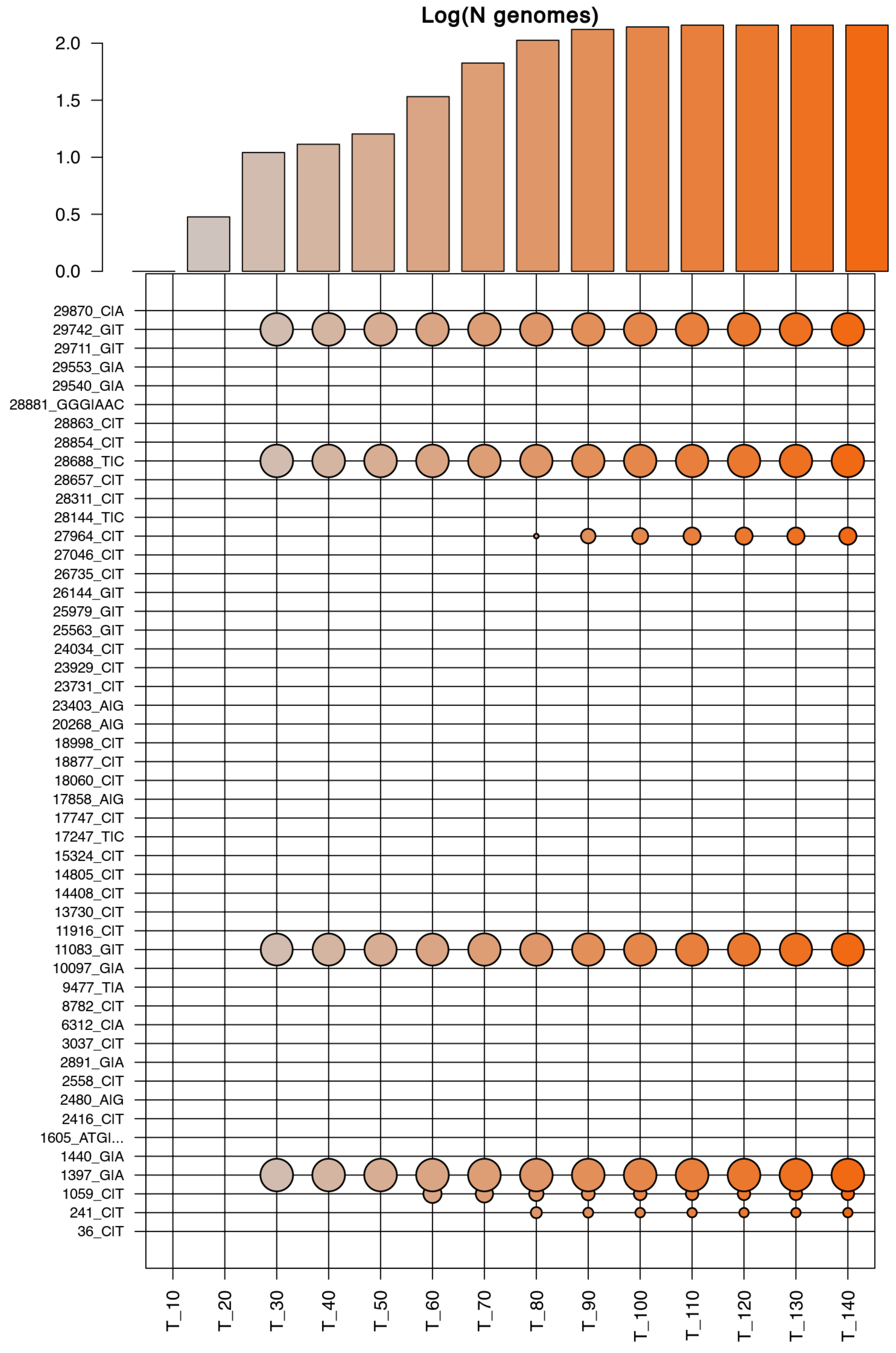


C11

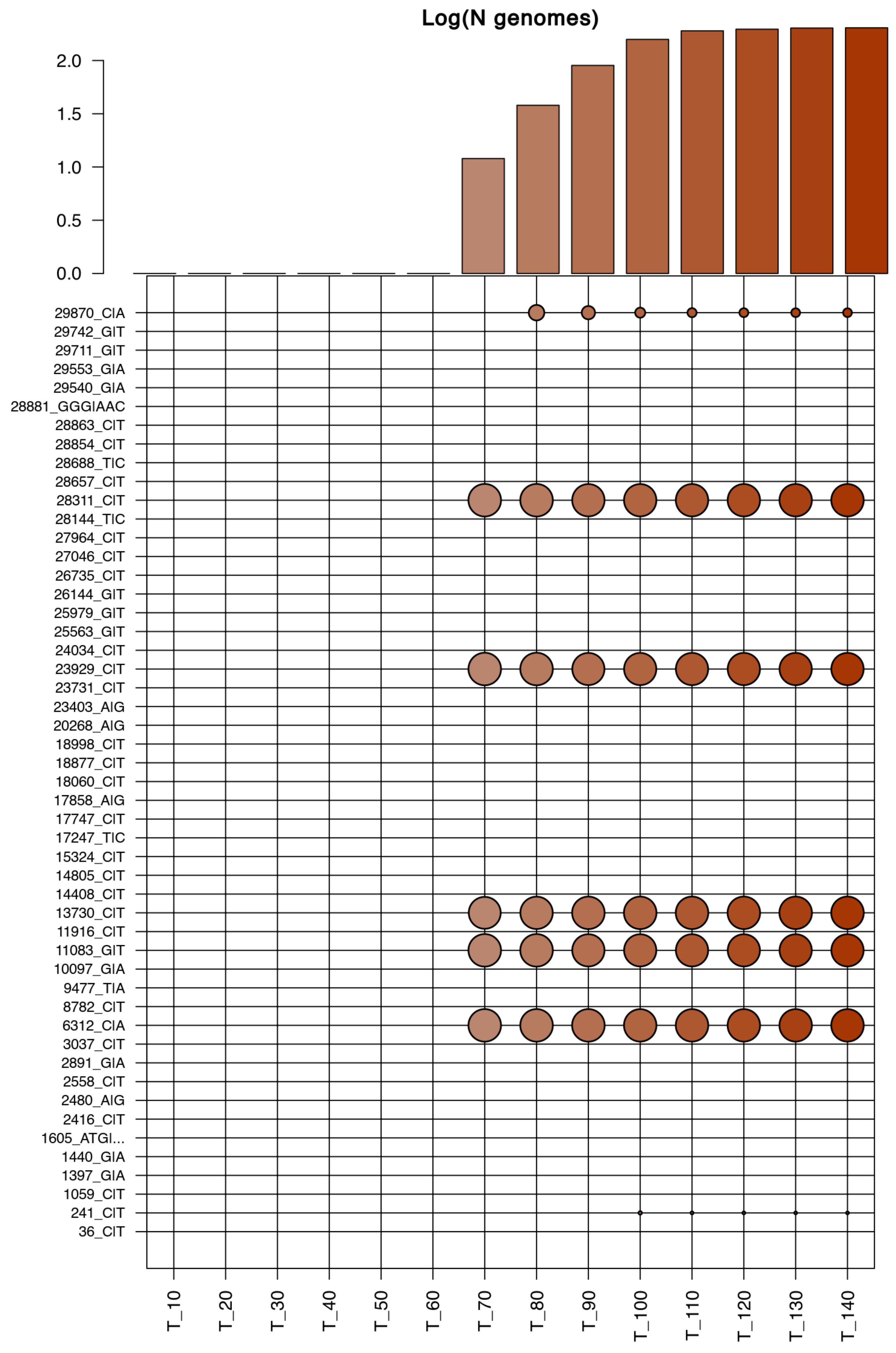


C12

### Supplementary Figure S8

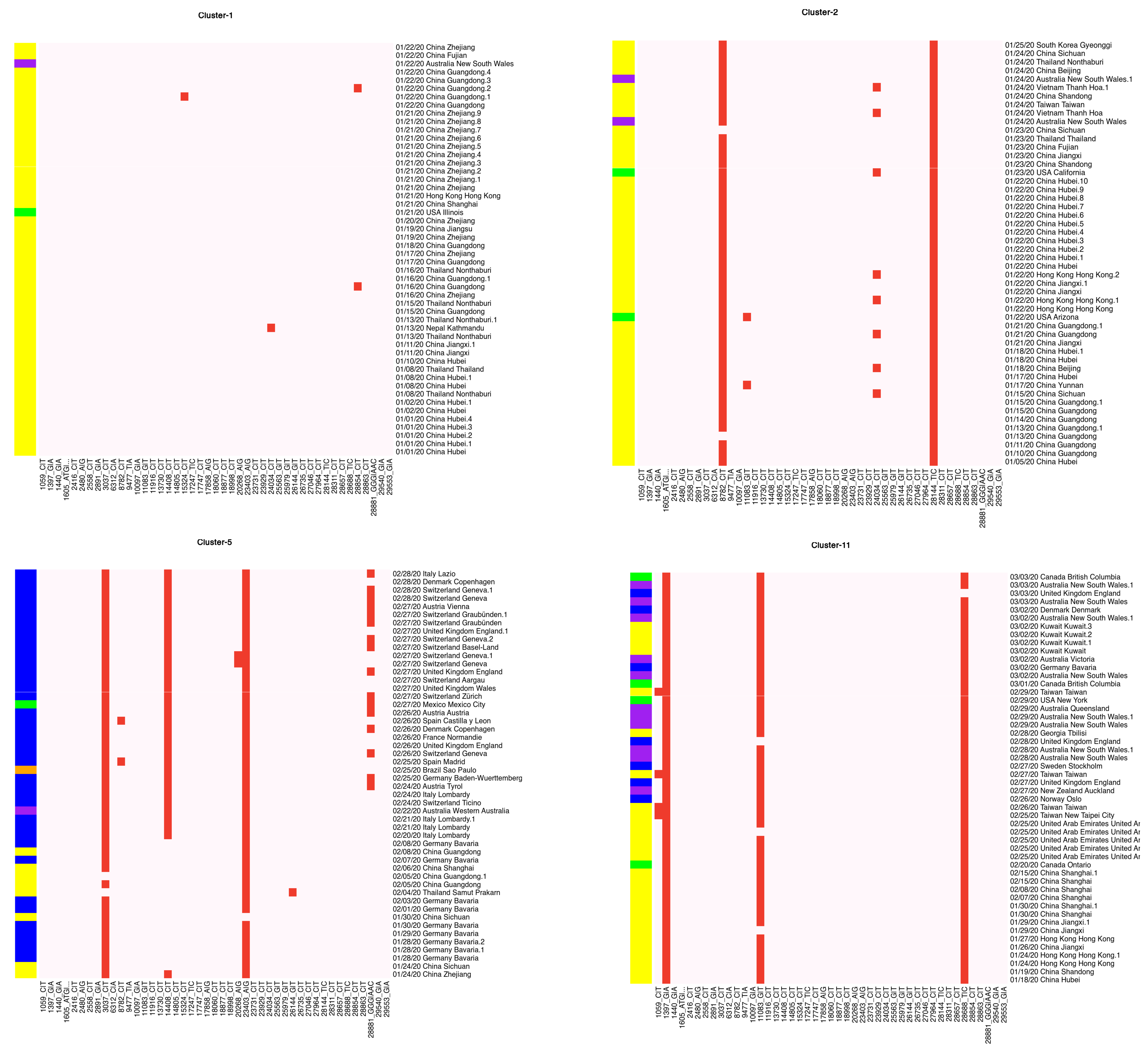

### Supplementary Figure S9

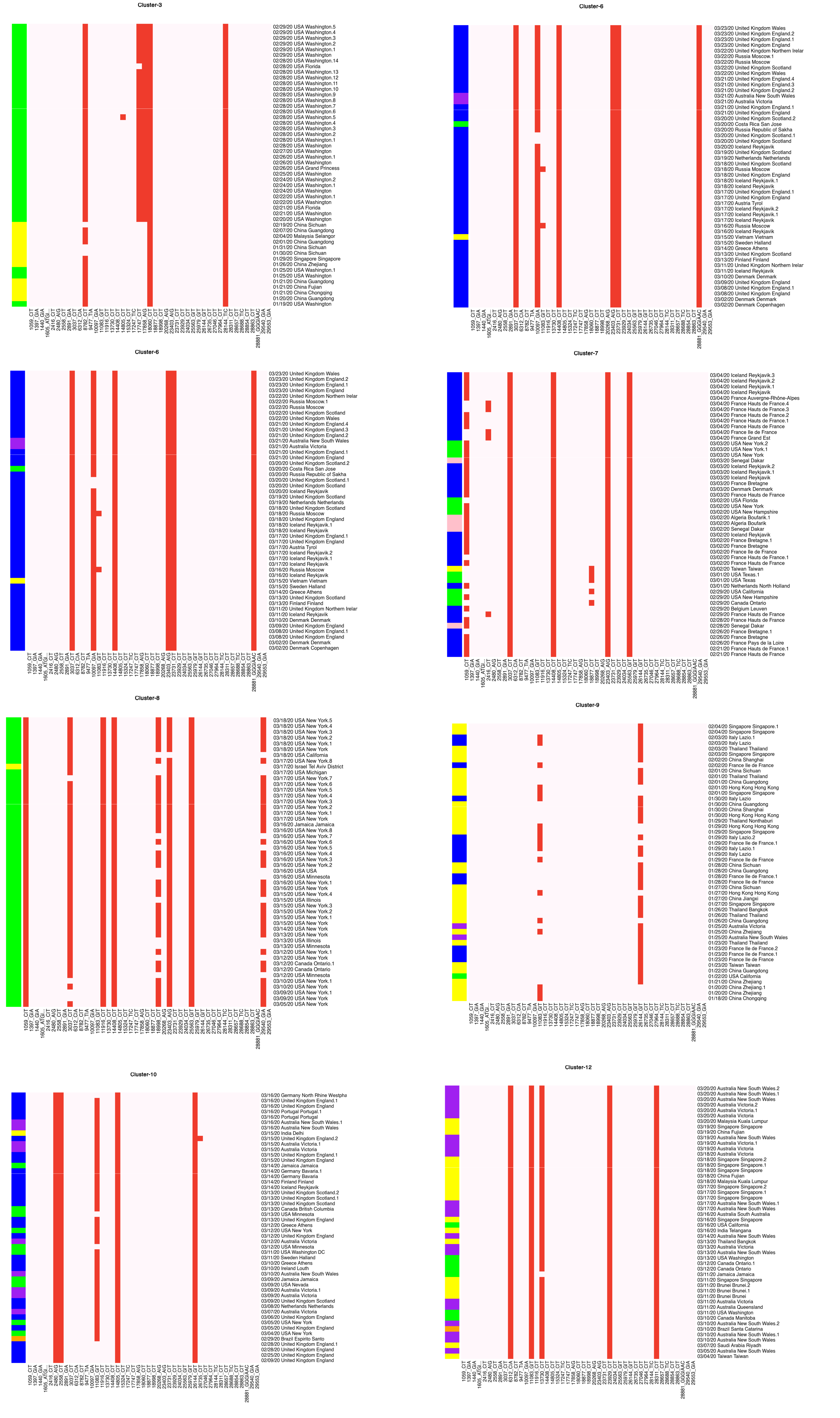

### Supplementary Figure S10

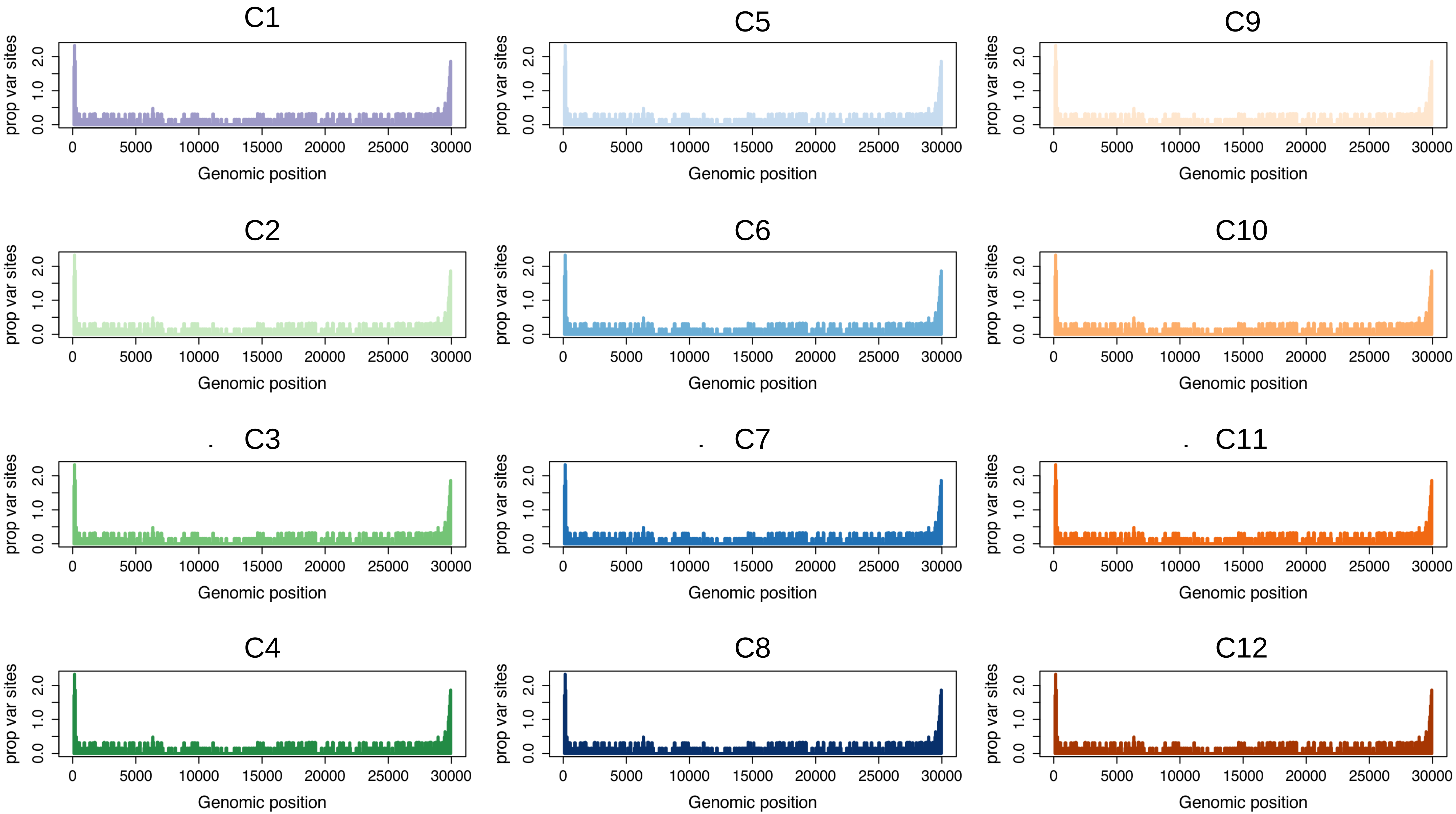
